# Mapping trait versus species turnover reveals spatiotemporal variation in functional redundancy in a plant-pollinator network

**DOI:** 10.1101/2021.11.29.470359

**Authors:** Aoife Cantwell-Jones, Keith Larson, Alan Ward, Olivia K. Bates, Tara Cox, Frida Brannlund, Charlotte Gibbons, Ryan Richardson, Jason M. Tylianakis, Jacob Johansson, Richard J. Gill

## Abstract

Functional overlap between species (redundancy) shapes competitive and mutualistic interactions, determining community responses to perturbations. Most studies view functional redundancy as static, even though individuals within species vary in traits over seasonal or spatial gradients. Consequently, we lack knowledge on trait turnover within species, how functional redundancy spatiotemporally varies, and when and where interaction networks are vulnerable to functional loss. Studying an Arctic bumblebee community, we investigated how body-size turnover over elevation and season shapes their host-plant interactions, and test how sensitive networks are to sequentially losing body-size groups. With trait turnover being larger than species, we found: i) late-season networks were less specialised when nodes comprised functionally similar bumblebees; ii) removal of bumblebee-body-size groups over species accelerated coextinction of host plants, with the magnitude varying in space and time. We demonstrate functional redundancy can vary spatiotemporally, and functional loss impacts interaction partners more than expected from species loss alone.

## 1. INTRODUCTION

Understanding the processes structuring communities over environmental gradients is central to determining how they will respond to environmental change (Pellissier *et al*., 2018; Orr *et al*., 2021). In particular, our ability to predict how organisms interact through trophic, mutualist or competitive processes depends on being able to map how functional traits vary in space and time, both within and between species (Gray *et al*., 2018; Gravel *et al*., 2019; Gómez *et al*., 2020; Hurtado *et al*., 2020). While key life-history traits directly determine inter- or conspecific relationships and the distribution of individuals, such traits are also simultaneously being selected and filtered by the environment (Gillespie *et al*., 2017; Bladon *et al*., 2020). Consequently, the frequency distributions of traits within populations can vary across environmental gradients and over seasons (Jourdan *et al*., 2016; Boyle *et al*., 2020; Taylor-Cox *et al*., 2020; Gutiérrez & Wilson, 2021). Yet how population-level responses combine to influence community trait-composition, and subsequently how spatiotemporal trait turnover feeds back to shape community interactions, remain poorly understood (Ings *et al*., 2009; Tylianakis & Morris, 2017; Peralta *et al*., 2020).

For insect pollinators, how functional traits are spatiotemporally distributed likely determines the occurrence of interactions with mutualistic host plants (Bewick *et al*., 2013; Brosi & Briggs, 2013), with compatibility between pollinator traits and plant hosts mediating interaction strengths (Bartomeus *et al*., 2016; Peralta *et al*., 2020). Investigating the dynamics of plant-pollinator relationships, however, has often been based on network comparisons between sites separated by large geographic distances or between years (Dupont *et al*., 2009; Cirtwill *et al*., 2018; Bascompte *et al*., 2019). This is despite individual interactions likely happening over distances of typically just hundreds of meters and over weekly (even daily) timeframes. Consequently, our understanding of plant-pollinator network lability at localised geographical scales and over short time frames remains limited. We must thus move beyond viewing plant-pollinator networks as geographically or seasonally static entities (Trøjelsgaard & Olesen, 2016; Tylianakis & Morris, 2017; Bramon Mora et al., 2020), and understand how species versus functional traits spatiotemporally turn over in localised communities (Robroek *et al*., 2017). In doing so, we can resolve the contribution of functional traits to network structure and robustness to perturbation. With trait variation responding to many environmental factors, studies mapping the distribution of trait variability can more accurately reveal how environmental change can shape and potentially threaten plant-pollinator interactions.

Bipartite networks can investigate how pollinator communities interact with their host plants by aggregating individuals into nodes (Ings *et al*., 2009; Poisot, Stouffer & Kéfi, 2016). In the past, nodes typically represented taxonomic units (Bascompte & Jordano, 2007), but more recently functional traits have been incorporated into networks by assigning each species (node) a mean value of a given trait from a set of sampled individuals (e.g. Bartomeus, 2013; Coux *et al*., 2016; Dehling *et al*., 2016). This assumes however that all individuals within each node are functionally equivalent and overlap in trait values between species is low – which is rarely the case (Gentile *et al*., 2021). Network structure changes markedly when individuals rather than species are treated as nodes (Tur *et al*., 2014), with intraspecific trait variation potentially being as large as interspecific (Des Roches *et al*., 2018), arising through sexual dimorphism (Bolnick *et al*., 2011), conditions during development (Fenberg *et al*., 2016) and adaptation to environmental gradients (Hurtado *et al*., 2020; Taylor-Cox *et al*., 2020). Additionally, temporal intraspecific variation can result from phenological variation with members of a species emerging at different times (Dongmo *et al*., 2018; Fric & Konvicka, 2002). Thus, to assess how functional trait turnover determines plant-pollinator network dynamics, we must scale down to the functional traits of individuals within populations, to disentangle how traits shape networks (e.g. Dupont *et al*., 2014; Kuppler *et al*., 2016; Rumeu *et al*., 2018). To our knowledge, however, data on individual traits and interactions at such fine spatiotemporal scales are generally lacking. Moreover, there are few empirical studies that incorporate intraspecific trait variation when studying functional contribution to plant-pollinator network structure (but see Dupont *et al*., 2014; Classen *et al*., 2017).

Here, we explore how individual variability in traits and interactions contribute to spatiotemporal dynamics in pollination network structure along an environmental gradient. We hypothesise: (1) intraspecific trait variability will be sufficiently large that functional turnover in space and time will exceed species turnover; (2) interaction networks where pollinator individuals are grouped together into nodes based on their traits, rather than species identity, will better capture interacting guilds of individuals. Subsequently, functional nodes will interact with a limited set of trait-matched plants to generate more-modular and less-connected networks, relative to taxonomic nodes; (3) loss of pollinator functional nodes will result in greater extinction cascades of host plants than species nodes. Addressing this latter hypothesis will test the degree to which individual functional trait redundancy can provide network robustness to pollinator species loss, and how dependent this is in space and time.

We test these hypotheses in a montane Arctic community of bumblebees, using bumblebee-plant visitation data collected repeatedly along an elevational transect (33 days) over a 57-day period. This temporal and elevational cline allowed us to map plant-bumblebee interaction and trait turnover in space and time. We focus on body size, as it is influenced by thermal conditions during development, and determines foraging capabilities at low and high relative temperatures (Lundberg & Ranta, 1980; Greenleaf *et al*., 2007). Body size also strongly correlates with proboscis length (influencing floral compatibility; Cariveau *et al*., 2016) and can predict pollinator-plant interactions (Eklöf *et al*., 2013; Klumpers *et al*., 2019). Additionally, as with many functionally important species, bumblebee body size at the population level does not remain constant through the season, as earlier emerging individuals have larger bodies (queens) than later emerging individuals (workers).

## 2. METHODS

### 2.1 Study site

We collected data along an elevational transect (420-1164 m.a.s.l.; est. 1917; Fries, 1925; MacDougall *et al*., 2021) on the eastern slope of Mt Nuolja in Abisko National Park, Sweden (68° 22’ N, 18° 47’ E). Along the 3.4 km transect, there are 13 permanent plots measuring 45 × 45 m, within which we recorded bumblebee activity (Fig. 1 & S1; SI 1). Five relatively distinct vegetational zones (Fig. 1 & S2; Table S1) are found along this transect and following Lundberg & Ranta (1980) can be described as: A) old Birch forest (420-550 m; *n* = 3 plots); B) young, newly established Birch forest (550-650 m; *n* = 2); C) shrub Willow (650-850 m; *n* = 3); D) herbaceous meadow (850-1050 m; *n* = 2); E) Arctic dwarf shrub and heath (1050-1164 m; n = 3). The number of plots (*n* = 13) was determined by the feasibility under which sampling could take place, dependent on both the terrain and number of observations that could be achieved in one day.

**Figure 1:**
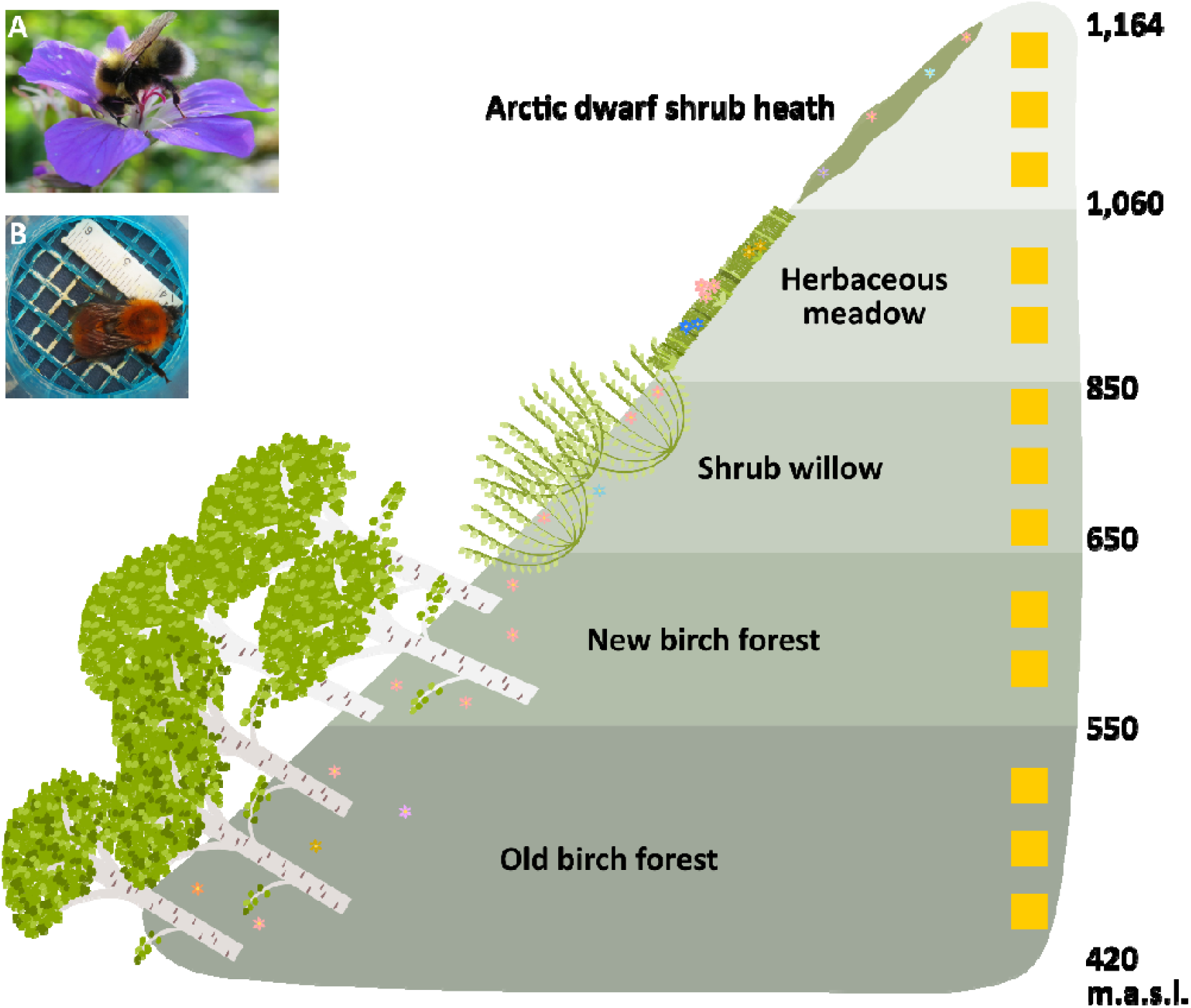
The elevational transect on Mount Nuolja spans five vegetation zones with 13 sampling plots (yellow squares). Diagram not to scale. Inset images show: (A) *Bombus jonellus* foraging on *Geranium sylvaticum*; (B) *B. pascuorum* inside a marking cage for measuring the intertegular distance (photo credits: Olivia Bates, Tara Cox & Aoife Cantwell-Jones).

### 2.2 Field observations and body size measurements

We conducted observations between 24^th^ May to 20^th^ July 2018, weather permitting. We typically conducted a “survey” – defined as undertaking bumblebee observations for all 13 plots over a two-day period (intended to be two consecutive days) – twice per week (Table S1). We observed each plot for a standardised 20 minutes between 08:00 - 19:00. For each survey, we randomised the time of day the plot was sampled (SI 2 for further details).

We recorded all bumblebees observed flying through or foraging inside the plot (*n* = 1,582), but only included those that were confidently identified taxonomically in the analyses (13 species; *n* = 1,047 individuals). If the bee was foraging on a flower, we recorded the plant species (*n* = 31; Table S2). We attempted to catch all observed bees with a butterfly net (45 cm diameter), and, if successful, transferred them (*n* = 709) into separate lidded, plastic holding pots (150 mL volume) and placed them in a dark insulated bag. At the end of the observation period, we transferred each bee in order of capture to a marking cage and took a top-down dorsal image using a digital camera (Canon SX720 HS) once the bee was still (e.g. Fig 1, inset B). From the images, we: i) confirmed species identification (consistent with Söderström (2017)), although *Bombus polaris* and *B. alpinus* were indistinguishable (Williams *et al*., 2015) and therefore grouped as a species aggregate; ii) measured the intertegular distance (ITD; distance between left and right forewing attachment points, which is an accurate proxy for body mass (Cane, 1987) and proboscis length (Cariveau *et al*., 2016)), by digitally landmarking each tegulum using TPS software (tpsDig v.2.3.2 & tpsUtil v. 3.2; Rohlf, 2015) and exporting the landmarks to the geomorph package in R to calculate scaled distance in millimetres (Adams *et al*., 2020). Of the bees with ITD measurements, we observed 69.8% (n = 495) foraging on a flower, and these comprised 94 queens, 270 workers, 21 drones, and 110 females where caste could not be confidently assigned (Table S3-5; SI 3).

### 2.3 Assessing intra- and interspecific variation in bumblebee body size

To gauge the magnitude of intraspecific variation in body size, we quantified the degree of overlap in ITD distributions between species by calculating the “coefficient of overlapping” (Ridout & Linkie, 2009) of pairwise bumblebee species kernel density estimates using the *overlapTrue* function of the “overlap” package (Meredith & Ridout, 2020). Output values range between zero and one, with one representing complete ITD distribution overlap between species. We only included species with ≥10 individuals that had ITD measured, yielding 45 species pairwise combinations. Additionally, we performed a mixed-effects ANOVA to test whether bumblebee species identity could significantly predict mean differences in ITD. Caste was nested within species included as random intercepts, to ensure that differences between species were not due to one species containing more of a given caste.

To analyse how ITD varied in space and time at the community level, we first fitted ordinary least squares models, with time expressed as calendar date, and altitude as the mean elevation of each sampling plot. Then, to assess if species responded similarly across time and space, we fitted mixed-effects models (using the “lme4” package; Bates *et al*., 2018) and included species as random intercepts with random slopes (either calendar date or mean elevation, respectively). We used likelihood ratio tests (LRT) to determine whether including random intercepts or slopes improved model fit.

### 2.5 Assigning individuals to functional groups

We created individual trait-based functional groups following the method of Rumeu et al. (2018), by aggregating individual bees based upon similarities in ITD and ignoring species identity (using the “cluster” package; Maechler *et al*., 2014). When clustering, we only included flower-visiting bees that had ITD measurements (n = 495; as we use these clusters in later analyses of plant-bumblebee visitation networks) and performed it on the pooled set of observations across the spatiotemporal gradient. We “constrained” the number of functional groups to match the total number of species present, to enable comparisons without confounding effects of changing number of nodes (Rumeu *et al*., 2018). The resulting 13 functional groups varied in abundance and ITD range (Fig. S3; Table S6) and were numbered in order of increasing body size (i.e. Constrained (C) 01 contained the smallest-bodied individuals).

### 2.6 Assessing spatiotemporal changes in species and functional-trait composition

To investigate spatial variation in the bumblebee community, we grouped plots from vegetation types A and B to represent a “low-elevation” dataset, and plots from vegetation types C, D and E, a “high-elevation” dataset. Pooling data across multiple plots was done to ensure sufficient sample size for our analyses across each point in space and time. Within each spatially pooled set of observations, we split observations into an “early” and “late” part of the season. To guide this split we considered colony-lifecycle phenology of the social bumblebees (queens emerge and dominate during the early season and workers during the late) to be natural stages of comparison to look at temporal functional turnover and determined the cut-off between “early” and “late” as occurring when 50% of observed female bees were likely to be queens (23^rd^ June 2018; Table S7; Fig. S4; SI 4). We estimated spatiotemporal turnover in the taxonomic and functional composition of communities using Bray-Curtis distance matrices (using the “betapart” package; Baselga *et al*., 2018), which are robust to under-sampling and taxonomic misidentification (Schroeder & Jenkins, 2018). We also decomposed the Bray-Curtis distances into their two additive components (sensu Baselga, 2013). Bray-Curtis distances range from zero to one, with one representing complete community turnover. To ensure the robustness of our results, we additionally computed Jaccard distances with a presence-absence version of the community data (using “vegan” package; Oksanen *et al*., 2007), the results of which can be found in the supplementary.

### 2.7 Comparing network architecture between species- and functional-group-based networks

We constructed two types of weighted networks from the same observed interaction data using the “bipartite” package (Dormann *et al*., 2009): 1) using species identity as nodes, 2) using functional group identity as nodes (see 2.5). We described each network using: the number of bumblebee nodes (taxonomic or functional groups) (B) and plant species (P); and the total number of observed interactions (I). To investigate whether the functional-group-based networks better capture trait matching, we additionally calculated: *Weighted connectance* (herein, connectance; e.g. Kaiser-Bunbury *et al*., 2011) to reflect the proportion of realised links in the network out of all possible links (i.e. 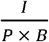, ranging between zero (unconnected network, suggesting many specialised interactions) and one (completely connected network); and *Weighted quantitative modularity Q* (herein, modularity; Beckett, 2016) to describe the presence of distinct, highly connected subnetworks. Modularity was calculated using the DIRTLPA+ algorithm (Beckett, 2016) and ranges from zero (the network does not have more links within subnetworks – modules – than expected by chance) to one (all links are within modules; highly modular network).

To assess whether the observed interaction networks were significantly less connected or more modular than expected, we standardised raw values using the mean and standard deviation of indices calculated from 999 null networks (z-scores). We generated these null networks using the *vaznull algorithm* (Vázquez *et al*., 2007) from the “bipartite” package (Dormann *et al*., 2009). These z-scores enabled us to estimate the likelihood, as *p*-values, of the observed network architectures emerging from random associations.

### 2.8 Using stepwise extinctions to compare network robustness between species- and functional-group-based networks

Through a randomised removal and iteration process we investigated the relative contribution of taxonomic and functional loss towards the rate of network collapse, based on the principle that interaction losses lead to subsequent host-plant extinctions. Here, an extinction occurred when a node had zero links remaining, with the assumption that no interaction rewiring occurred after extinction. By comparing the differential rate of network collapse under functional vs. taxonomic loss, we can investigate which poses a greater threat to plant-pollinator interactions under environmental change, and whether this depends on the spatiotemporal context. We utilised a novel method of stepwise extinction that focuses on the frequency distribution of ITD within species. We removed one discrete portion of a node, where, for the species-based networks, we removed a functional group from a node (e.g. all individuals with ITD 4.04-4.52 mm being removed from *B. jonellus*), and, for the functional-group-based networks, we removed a species from a node (e.g. *B. alpinus/polaris* was removed from the node containing individuals with 7.71-7.95 mm ITDs). We performed this removal randomly until one of the trophic levels (plant or bumblebee) had only one node remaining. For example, during the late season at low elevation, there are 37 unique species × functional group combinations (number of functional groups within species, or vice versa), meaning a maximum of 37 removals could occur. After each removal, we measured the number of remaining flower species, (weighted) connectance and (weighted quantitative) modularity – we used the rate of change in these metrics as proxies of network robustness. We iterated the process of randomised removal and calculation of indices 1,000 times and performed it separately for each point in space and time.

Using the results from the iterations, we modelled the change in each measure of network robustness as a function of the number of “items” (viz. discrete portions within nodes) that had been removed from each network, using generalised least squares models (gls; from “nlme” package; Pinheiro *et al*., 2017). Models took either linear or quadratic form, depending on the R^2^ value. To account for increasing variance of model residuals when items were removed, we included an exponent-of-the-variance covariate. Due to issues with model convergence, we were unable to account for increasing model-residual variance for modularity, so these results have been placed in the supplementary materials, and should be interpreted with caution. To assess and compare levels of “network robustness”, we looked at two aspects: i) differences in the shape of the plotted line estimates; ii) area under each curve (AUC) per model (“DescTools” package; Signorell, 2021). We standardised by dividing the raw AUC value by the product of the number of items removed (as some points in space and time contained more functional groups and species) and the maximum value of the calculated index. For the number of remaining plant species, if loss of functional groups has a larger effect than loss of species, we should expect functional-group networks to show lower standardised AUC, as it indicates that fewer individuals have to be removed from the network before its structure collapses. Alternatively, for weighted connectance, a higher AUC when removing functional groups would suggest the network requires fewer individuals to be removed for specialised interactions to be lost (i.e. before it becomes fully connected).

We performed all analyses using R (v. 4.0.2; R Core Team, 2020).

## 3. RESULTS

### 3.1 How do functional groups and species turn over in space and time?

Bee species identity did not significantly predict ITD (mixed-effects ANOVA: *F* = 0.36, *p* = 0.97; Table S8), with the 13 species showing considerable overlap in ITD distributions (Fig. S5). While bumblebee ITD ranged from 3.21 to 9.43 mm, 80% of the pairwise species comparisons (n = 45) showed overlapping ITD kernel density estimates of ≥0.5, and 11% showed ≥0.75 (Table S9). *B. jonellus and B. pratorum*, however, frequently deviated from this trend, each constituting four (of nine) pairwise comparisons with <0.5 overlap.

At the community level, mean ITD significantly decreased over the season (lme: -0.0491 mm day^- 1^ ± 0.00251, *t* = -19.6, *p* <0.001; *N* = 709; Fig. 2; Table S10-12) and was consistent across bumblebee species (likelihood ratio tests (LRT) supporting species having random intercepts (LRT: *p* <0.001), but not random slopes (LRT: *p* = 1) for the effect of season). ITD also significantly increased with elevation (lm: 0.121 mm 100 m^-1^ ± 0.0279, *t* = 4.33, *p* <0.001; *N* = 709; Fig. 2; Table S13), with species having different intercepts (LRT: p <0.001; Table S14-15) but similar slopes (LRT: *p* = 0.195). 3.1.2

**Figure 2:**
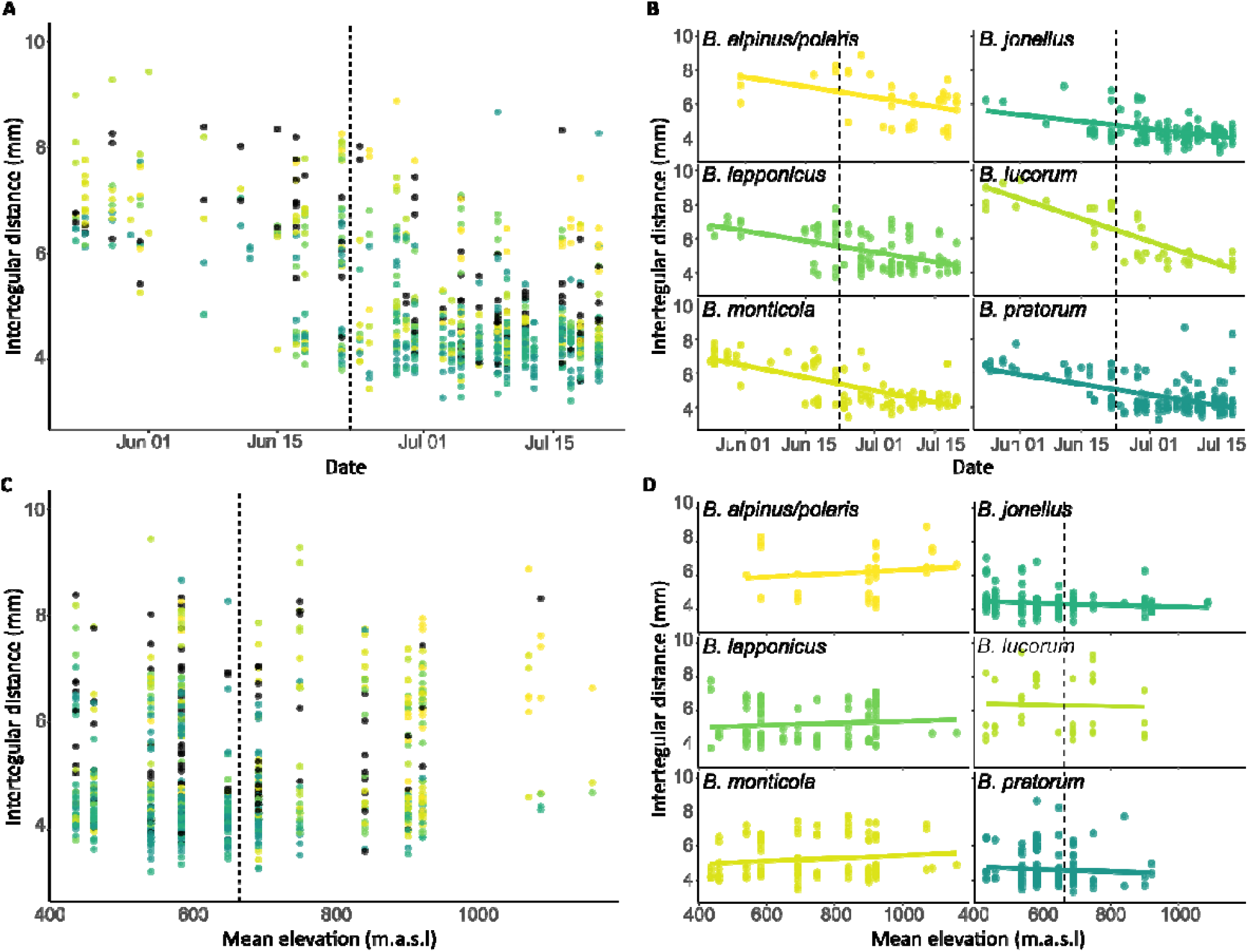
Body size (ITD) turnover in space and time for the bumblebee community. Each data point represents a captured individual. Panels A-B show ITD variation as the season progressed at A) the community level and B) for the six most abundant species, with the dashed line depicting the transition from queen-to worker-dominated foraging, based on estimates from a binomial generalised linear model (Table S7). Panels C-D show ITD variation along the elevational gradient at C) the community level and D) for the six most abundant species, with the dashed line depicting the split between assigned low-(left of line) and high-(right) elevation survey plots. The six most abundant species (*Bombus alpinus/polaris, B. jonellus, B. lapponicus, B. lucorum, B. monticola and B. pratorum*) are colour coded, with the colours in A & C matching those in B & D. The remaining seven species (*B. balteatus, B. bohemicus, B. cingulatus, B. flavidus, B. hortorum, B. hyperboreus and B. pascuorum*) are black in A & C.

#### 3.1.2 Changes in species versus functional composition in the community

Species turnover was similar in time and space (Bray-Curtis distance (D_BC_): 0.474-0.672 & 0.322-0.781, respectively; Fig. 3; see Table S16-18 for breakdown in the additive components of D_BC_ and Jaccard dissimilarity). Functional turnover, however, was larger than species turnover (mean D_BC_ ± standard error: 0.651 ± 0.099 vs. 0.537 ± 0.068, respectively; Table S17). As suggested by 3.1.1 and Fig. 2, functional turnover was also greater over the season than with elevation (D_BC_: 0.776-0.792 & 0.269-0.500, respectively). Over the season, we saw an increased relative abundance of functional groups containing smaller individuals: C02 (mean ITD = 3.82 mm), C03 (4.27 mm) and C04 (4.71 mm), and decrease in the larger-bodied functional groups: C07 (6.23 mm) and C08 (6.69 mm). With elevation, groups C09 (7.22 mm) and C10 (7.81 mm) increased in relative abundance, while instead C03 decreased.

**Figure 3:**
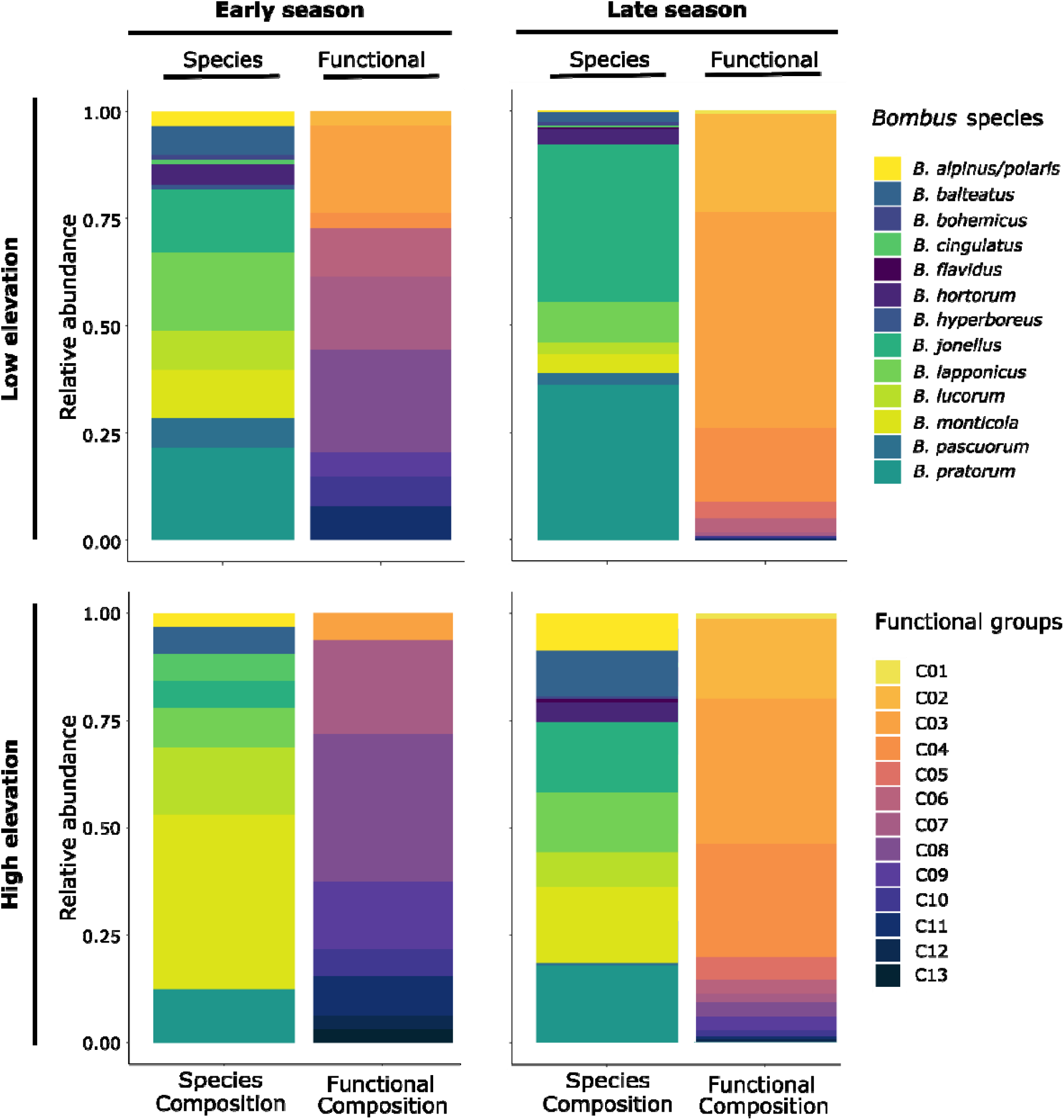
Turnover of species- and functional-group-based communities in space (Low & High elevation) and time (Early & Late season). The number of individuals in the species- and functional-group-based bumblebee communities are the same (early season, low elevation = 88; late, low = 224; early, high = 32; late, high = 151). Functional groups are numbered from the smallest-bodied individuals (C01) to the largest (C13).

### 3.2 Differences in architecture between species and functional networks over space and time

Connectance and modularity differed in the two network types across space and time (Fig. 4), with networks in the late season being significantly less connected and more modular than expected under the null distribution (species (s): late season, low elevation connectance: *z* = -3.82, *p* = <0.001, modularity: *z* = 5.17, *p* <0.001; late, high connectance: *z* = -2.02, *p* = 0.043, modularity = *z* = 4.98, *p* <0.001; functional (f): late, high connectance: *z* = -3.04, *p* = 0.002; Table S19). During the late season, species-based networks were also consistently more modular than functional-group-based networks (low: s = 0.241, f = 0.136; high: s = 0.334, f = 0.245), although both showed similar levels of connectance (Table S19).

**Figure 4:**
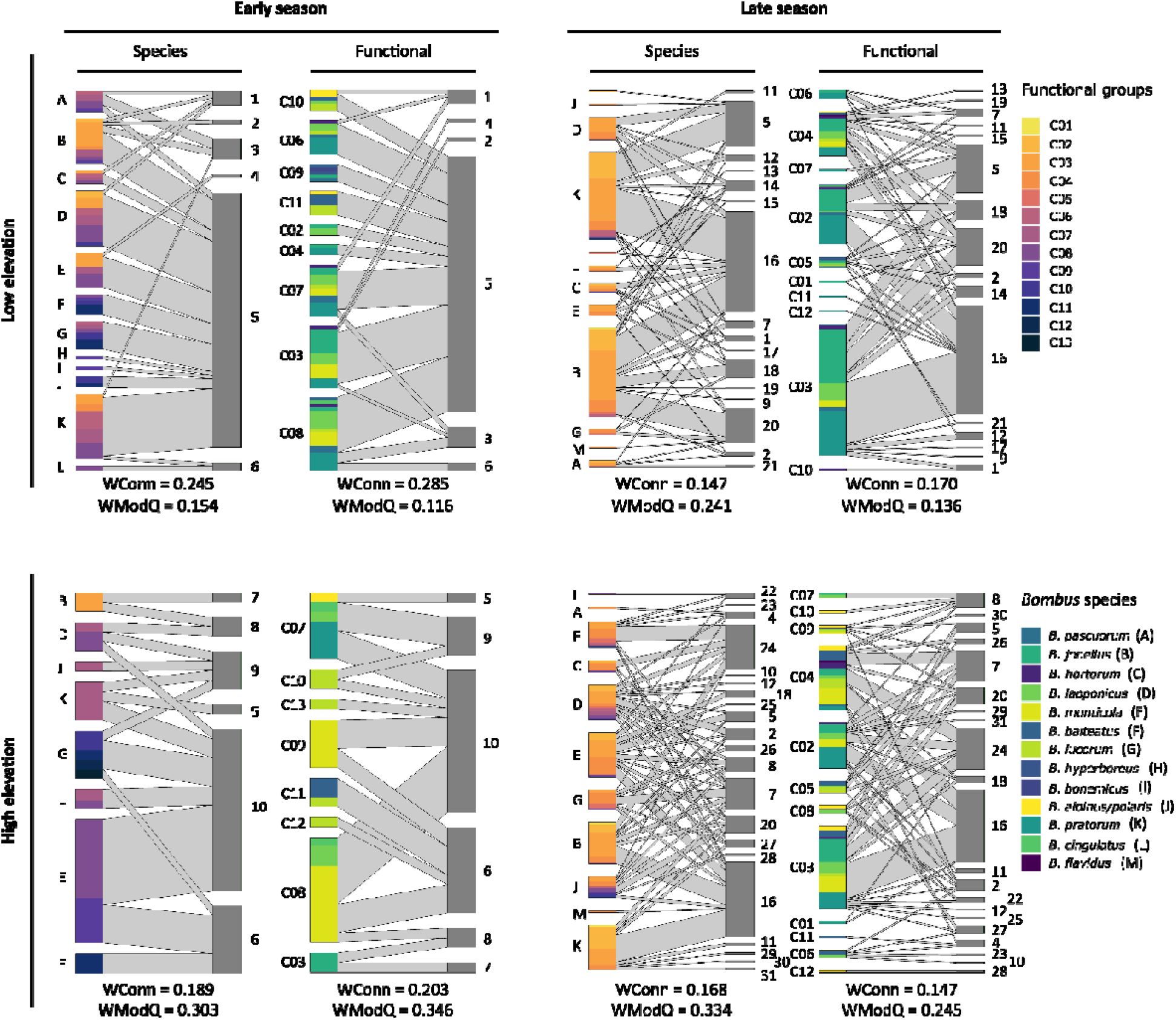
Species- and functional-group-based bumblebee-plant visitation networks at different points in space and time. Nodes in grey represent plant species: 1) *Andromeda polifolia*, 2) *Pedicularis lapponica*, 3) *Salix myrsinites*, 4) *Phyllodoce caerulea*, 5) *Vaccinium myrtillus*, 6) *Salix phylicifolia*, 7) *Trollius europaeus*, 8) *Salix glauca*, 9) *Salix myrsinifolia*, 10) *Salix lanata*, 11) *Melampyrum sylvaticum*, 12) *Solidago virgaurea*, 13) *Silene dioica*, 14) *Myosotis decumbens*, 15) *Rubus saxatilis*, 16) *Geranium sylvaticum*, 17) *Linnaea borealis*, 18) *Vaccinium uliginosum*, 19) Saxifraga aizoides, 20) *Vaccinium vitis-idaea*, 21) *Melampyrum pratense*, 22) *Taraxacum spp*., 23) *Ranunculus acris*, 24) *Astralagus alpinus*, 25) *Epilobium alsinifolium*, 26) *Pyrola rotundifolia*, 27) *Potentilla crantzii*, 28) *Diapensia lapponica*, 29) *Bartsia alpina*, 30) *Pyrola minor*, and 31) *Viola biflora*. For each bipartite network, the coloured nodes on the left trophic level represent bumblebee groups. For networks in the species column, we show the functional group composition of each species indicated by different colours (C01 representing the smallest-bodied bees and C13 the largest). For the networks in the functional column (where groups of bumblebees are clustered based on ITD similarity), we show the species composition of each functional-group node as indicated by different colours. At the bottom of each network is shown its raw weighted connectance (WConn) and weighted quantitative modularity (WModQ).

### 3.3 Differences in robustness between species and functional networks over space and time

For the stepwise removal analysis, removing functional groups from species nodes always led to faster extinction of flower species (f = 0.738-0.828, s = 0.765-0.838; Fig. 5A; Tables S20-21) than removal of bumblebee species. For flower species, this difference was most pronounced during the early season at high elevation, in which removing species from functional-group nodes showed a convex decline in number of flower species remaining, whereas removing functional groups from species nodes had a linear decline (AUCs: 0.800 vs. 0.738, respectively). For connectance, the differential rates at which the networks became fully connected under functional vs. taxonomic loss was spatiotemporally dependent. For example, at low elevation, and at high elevation during the early season, removing species from functional group nodes led to a faster loss of specialised interactions than removing functional groups from species nodes (i.e. higher standardised AUCs reflecting a more rapid increase in connectance; Fig. 5B; AUC range: 0.751-0.888 vs. 0.732-0.885, respectively). In contrast, at high elevation during the late season, this trend was reversed, with functional loss leading to a faster decline in specialised interactions (0.692 vs. 0.764, respectively).

**Figure 5:**
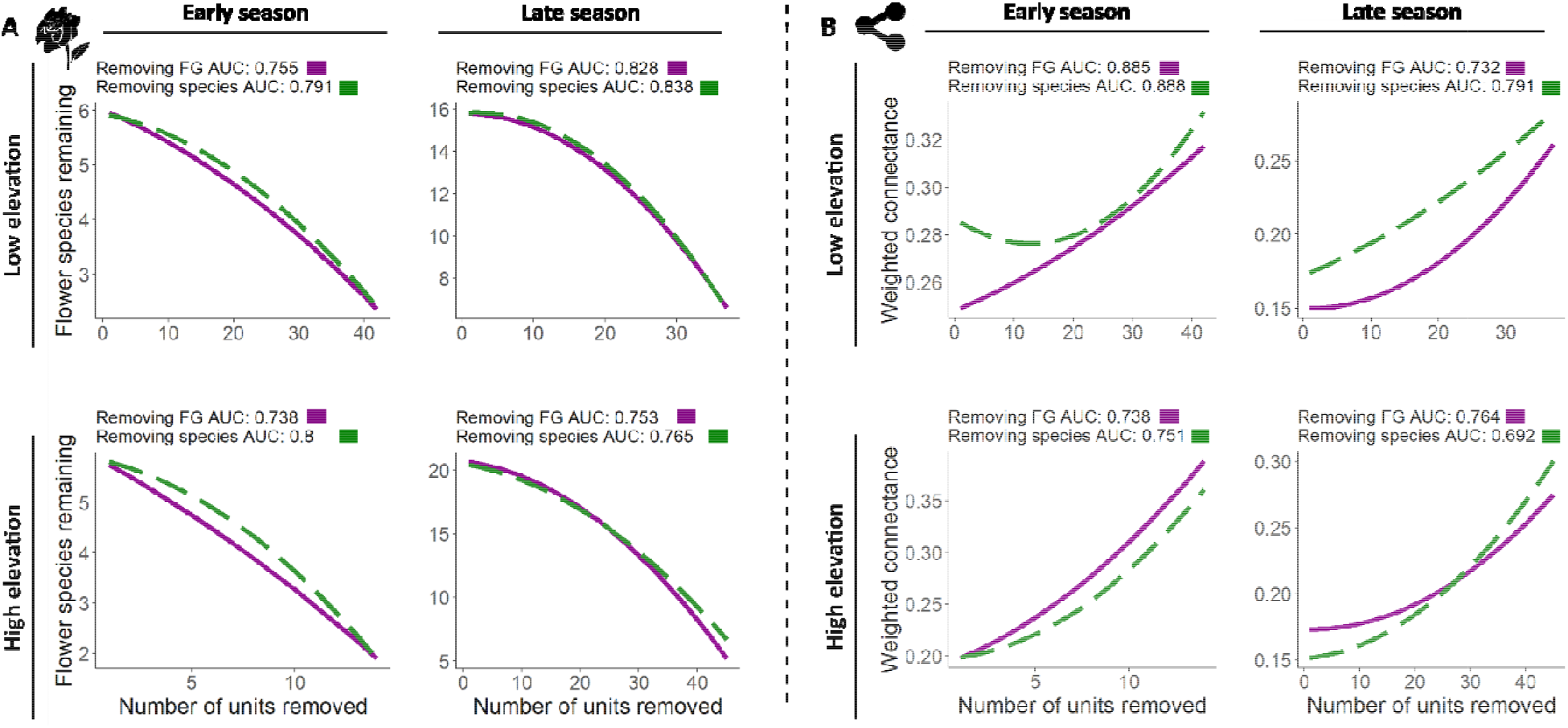
A) Number of remaining flower species and B) change in weighted connectance, in response to stepwise bumblebee removals to assess network robustness to interaction extinctions. Simulations randomly removed either species from functional-group nodes (green dashed line) or functional groups from species nodes (purple solid line), for 1,000 iterations. At each random removal, the number of remaining flower species and weighted connectance were calculated (for graphs with raw values, see Figs. S6-9). Standardised area under the curve (AUC) values are given for each plot. A) Higher AUC suggests slower co-extinction of flower species, i.e. greater robustness to extinction. B) Lower AUC suggests delayed loss of specialised interactions (longer time for network to become fully connected), i.e. greater robustness to extinction. FG = functional groups.

## 4. DISCUSSION

### 4.1 Mapping spatiotemporal trait change reveals decoupling of functional and species turnover

When considering the bumblebee community as a static entity, most bumblebee species overlapped in body size suggesting high functional redundancy. Species identity was thus a poor surrogate for individual trait variation. Additionally, body size within a species varied in space and time, such that functional turnover as the season progressed was far larger relative species turnover. While body size turnover in time is somewhat expected in bumblebees (large queens followed by smaller workers), we also found functional turnover in space, which again exceeded species turnover, particularly early in the season. This decoupling of species identities and their mean traits (Robroek *et al*., 2017) highlights why overlooking the spatiotemporal dynamics of networks, as well as assigning mean trait values to species, may poorly explain functional outcomes such as trait compatibility and resource acquisition (Rumeu *et al*., 2018; Wong & Carmona, 2021).

Spatial turnover being greater in functional than in taxonomic communities suggests that, while species adaptations may have contributed to higher altitude distributions (larger bodied species found at higher elevations, particularly in early season), intraspecific variation in body size still mediated where bees could be found. Smaller bees of any species were filtered from the community at high altitude (also see McCabe *et al*., 2019). This likely represents a plastic response at the colony level (Classen *et al*., 2017), where larger individuals within a species are able to forage at higher elevation, due to being better able to thermoregulate (Heinrich, 1975) and fly at lower temperatures (Kenna *et al*., 2021).

Temporal functional turnover being independent from species identity is not exclusive to bumblebees (Fukami *et al*., 2005; Classen *et al*., 2017; Robroek *et al*., 2017; Gómez *et al*., 2020), although the degree of such decoupling may vary across organisms. Yet, surprisingly little work has been done on mapping temporal trait distributions, especially at the spatiotemporal resolution or across a large community that we cover (but see Classen *et al*., 2017). Further, our study is one of the first to associate such a trait distribution with a key functional role – floral visitation as a proxy for pollination service (but see Ranta & Lundberg, 1981; Miller-Struttmann & Galen, 2014; Classen *et al*., 2017). Studies should therefore benefit from considering species as nested within a functional continuum, rather than as distinct units in terms of their traits (Bolnick *et al*., 2011; Siefert *et al*., 2015; Des Roches *et al*., 2018).

### 4.2 Architecture of functional interactions varied in space and time

Our plant-bumblebee interaction networks varied architecturally across space and time, contrasting with previously studied networks that showed relatively stable macroscopic features (e.g. connectance) across space (at local and regional scales) and over time (within seasons and across years; Olesen *et al*., 2008; Dupont *et al*., 2009; Trøjelsgaard & Olesen, 2016; Tylianakis & Morris, 2017). We found modularity to increase and connectance to decrease as the season progressed. This was associated with an increase in bumblebee abundance and plant species richness, which could have led to greater competition between bumblebee species promoting more specialised host-plant interactions (Brosi & Briggs, 2013; Miller-Struttmann & Galen, 2014). This is consistent with ecological character-displacement theory, in which competition leads to niche divergence in sympatric species; (Germain *et al*., 2018; Egas *et al*., 2005). Alternatively, other temporal changes in our studied plant-pollinator community in association with abiotic factors, like increasing mean daily temperatures, may have contributed. For example, Ohler et al. (2020) observed that microclimatic variation in temperature (of soils) was positively related to plant productivity and plant-visitor richness, and subsequently changes in network architecture. But ultimately, our findings underline how any static perspective of plant-pollinator interactions will miss nuanced but fundamental properties of how species spatiotemporally interact.

Functional over taxonomic networks being more connected and less modular later in the season was contrary to the expectation that functional networks would better represent trait compatibility between bumblebees and plants (Stang *et al*., 2009; Klumpers *et al*., 2019). This unexpected pattern could be due to the often generalised nature of pollination networks (Fort *et al*., 2016), meaning that trait matching might not always be strong. Inclusion of additional functional traits (Pigot *et al*., 2020), clustering of individuals based on their position in multidimensional trait space (Dehling *et al*., 2016; Coux *et al*., 2016), and/or measures of evolutionary history (Bascompte *et al*., 2019; Hurtado *et al*., 2020) might improve investigations of how trait compatibility determines the spatiotemporal dynamics of interactive networks.

### 4.3 The impact of functional group loss on network robustness is spatiotemporally dependent

When simulating bumblebee extinction, removing functional groups from networks led to a faster secondary coextinction of plant species than removing species, especially during the early season at high elevation. This is explained by our functional networks being more connected and less modular, enabling perturbations (species loss) to spread quickly (Kortsch *et al*., 2015). Functional changes to communities, such as declines in body size, have been shown to accompany environmental change drivers (e.g. Guthrie, 2003; Rode *et al*., 2010; Wu *et al*., 2019). Our results suggest that functional group loss may cause faster extinction of interaction partners than based on species loss alone especially given body size overlap was as large within as it was between species. It is plausible to lose whole functional groups, for example, if an early drought disproportionately affected larger bumblebees through reduced floral resources (Couvillon & Dornhaus, 2010) or if a late frost disproportionately affected smaller bees (Heinrich, 1975; Kenna *et al*., 2021). Studies focusing solely on species loss when investigating how communities may be affected by environmental change (e.g. Burkle *et al*., 2013; Bascompte *et al*., 2019; Soroye *et al*., 2020) may thus underestimate the vulnerability of communities to perturbations.

Bumblebee species removal accelerated the loss of specialised bumblebee-plant interactions relative to functional removal, except during the late season at high elevation, where the reverse occurred. These results highlight the importance of understanding how redundancy varies in time and space. Similarly, Wardle & Zackrisson (2005) found that empirical removal of species or functional groups affected ecosystem-level properties of some islands (in northern Sweden) but not others, depending on historical disturbance regime, island size and successional stage. The spatiotemporal context dependency of community interactions is thus important to consider when understanding network robustness (Thuiller *et al*., 2014; Robroek *et al*., 2017).

Our stepwise modelling approach assumed bumblebee group extinctions could occur randomly. Drivers of extinctions however likely occur and act non-randomly, such as land-use and climate change (Larsen *et al*., 2005), and thus informing extinction order probabilities will give new insights into rates of co-extinctions. We also assumed bumblebee-plant interactions did not rewire. Although commonly assumed in stepwise-extinction approaches (e.g. Classen *et al*., 2020; Keyes *et al*., 2021), this is unlikely in real communities (Brosi & Briggs, 2013; Bramon Mora *et al*., 2020). However, the lack of plant functional-trait data precluded predictions about which individuals would be sufficiently compatible in their traits to predict rewiring. Additionally, we treated each point in space and time in our extinction analysis as independent and species as independent from functional groups. This overlooks possible temporal autocorrelation in bumblebee biology (i.e. removing (large) queens early in the season could result in fewer (small or large) workers later), and does not consider that relatively large-bodied species could be more negatively affected if a large-bodied functional groups were lost (and vice versa for small-bodied species). To consider these in future analyses, we urge studies to collect trait and interaction data at fine spatiotemporal scales to elucidate mutualist and competitive responses to environmental change.

## 5. CONCLUSION

Our study reinforces why individual traits should be used to construct interaction networks, and the fundamental importance of where and when networks are studied during a season. The impact of functional or taxonomic losses on network robustness was not uniform in space and time, with the rates of plant coextinctions and loss of specialised interactions varying. However, losing bumblebee functional groups typically accelerated plant coextinctions more than losing species, meaning coextinctions of interaction partners could happen more quickly under functional loss than expected based on pollinator species extinction. Specifically for our study system, the loss of key functional groups dominating the early season and high altitude could be devastating for the whole plant-pollinator community. Overall, the level of redundancy within plant-pollinator community interactions and subsequently network robustness to perturbation is spatiotemporally dependent. Understanding how trait distributions change in localised space and time is thus essential to understanding dynamic population responses to environmental change.

## Acknowledgements

ACJ is funded by the NERC Science and Solutions for a Changing Planet doctoral training programme, Imperial College London. JMT is funded by the Marsden Fund (Grant number: UOC1705). The project was also supported by an INTERACT grant (funded by H2020 - agreement no. 730938) awarded to RJG. We would like to thank J. Gustafsson for help with plant identification, H. Rosenzweig for help setting up the sampling plots, and the Climate Impact Research Centre (CIRC, Umeå University) & Abisko Scientific Research Station (ANS) for field equipment and their continued support.

## REFERENCES

Adams, D.C., Collyer, M.L. & Kaliontzopoulou, A. (2020). Geomorph: Software for geometric morphometric analyses. R package version 3.2.1.

Bartomeus, I. (2013). Understanding Linkage Rules in Plant-Pollinator Networks by Using Hierarchical Models That Incorporate Pollinator Detectability and Plant Traits. PLoS One, 8, e69200.

Bartomeus, I., Gravel, D., Tylianakis, J.M., Aizen, M.A., Dickie, I.A. & Bernard-Verdier, M. (2016). A common framework for identifying linkage rules across different types of interactions. Funct. Ecol.

Bascompte, J., García, M.B., Ortega, R., Rezende, E.L. & Pironon, S. (2019). Mutualistic interactions reshuffle the effects of climate change on plants across the tree of life. Sci. Adv., 5, eaav2539.

Bascompte, J. & Jordano, P. (2007). Plant-Animal Mutualistic Networks: The Architecture of Biodiversity. Annu. Rev. Ecol. Evol. Syst., 38, 567–593.

Baselga, A. (2013). Separating the two components of abundance-based dissimilarity: balanced changes in abundance vs. abundance gradients. Methods Ecol. Evol., 4, 552–557.

Baselga, A., Orme, D., Villeger, S., De Bortoli, J., Leprieur, F. & Baselga, M.A. (2018). Package ‘betapart.’ Partitioning beta Divers. Turnover Nestedness Components. Ver, 1.

Bates, D., Maechler, M., Bolker, B., Walker, S., Christensen, R.H.B., Singmann, H., et al. (2018). Package ‘lme4.’ Version, 1, 437.

Beckett, S.J. (2016). Improved community detection in weighted bipartite networks. R. Soc. Open Sci., 3.

Bewick, S., Brosi, B.J. & Armsworth, P.R. (2013). Predicting the effect of competition on secondary plant extinctions in plant-pollinator networks. Oikos, 122, 1710–1719.

Bladon, A.J., Lewis, M., Bladon, E.K., Buckton, S.J., Corbett, S., Ewing, S.R., et al. (2020). How butterflies keep their cool: Physical and ecological traits influence thermoregulatory ability and population trends. J. Anim. Ecol., 89, 2440–2450.

Bolnick, D.I., Amarasekare, P., Araújo, M.S., Bürger, R., Levine, J.M., Novak, M., et al. (2011). Why intraspecific trait variation matters in community ecology. Trends Ecol. Evol.

Boyle, M.J.W., Bishop, T.R., Luke, S.H., van Breugel, M., Evans, T.A., Pfeifer, M., et al. (2020). Localised climate change defines ant communities in human-modified tropical landscapes. Funct. Ecol., 7, 14.

Bramon Mora, B., Shin, E., CaraDonna, P.J. & Stouffer, D.B. (2020). Untangling the seasonal dynamics of plant-pollinator communities. Nat. Commun., 11, 1–9.

Brosi, B.J. & Briggs, H.M. (2013). Single pollinator species losses reduce floral fidelity and plant reproductive function. Proc. Natl. Acad. Sci. U. S. A., 110, 13044–13048.

Burkle, L.A., Marlin, J.C. & Knight, T.M. (2013). Plant-pollinator interactions over 120 years: Loss of species, co-occurrence, and function. Science (80-.)., 340, 1611–1615.

Cane, J.H. (1987). Estimation of Bee Size Using Intertegular Span (Apoidea). J. Kansas Entomol. Soc., 60, 145–147.

CaraDonna, P.J., Petry, W.K., Brennan, R.M., Cunningham, J.L., Bronstein, J.L., Waser, N.M., et al. (2017). Interaction rewiring and the rapid turnover of plant-pollinator networks. Ecol. Lett., 20, 385–394.

CaraDonna, P.J. & Waser, N.M. (2020). Temporal flexibility in the structure of plant–pollinator interaction networks. Oikos, 129, 1369–1380.

Cariveau, D.P., Nayak, G.K., Bartomeus, I., Zientek, J., Ascher, J.S., Gibbs, J., et al. (2016). The Allometry of Bee Proboscis Length and Its Uses in Ecology. PLoS One, 11, e0151482.

Cirtwill, A.R., Roslin, T., Rasmussen, C., Olesen, J.M. & Stouffer, D.B. (2018). Between-year changes in community composition shape species’ roles in an Arctic plant-pollinator network. Oikos, 127, 1163–1176.

Classen, A., Steffan-Dewenter, I., Kindeketa, W.J. & Peters, M.K. (2017). Integrating intraspecific variation in community ecology unifies theories on body size shifts along climatic gradients. Funct. Ecol., 31, 768–777.

Couvillon, M.J. & Dornhaus, A. (2010). Small worker bumble bees (Bombus impatiens) are hardier against starvation than their larger sisters. Insectes Sociaux, 57, 193–197.

Coux, C., Rader, R., Bartomeus, I. & Tylianakis, J.M. (2016). Linking species functional roles to their network roles. Ecol. Lett., 19, 762–770.

Dehling, D.M., Jordano, P., Schaefer, H.M., Böhning-Gaese, K. & Schleuning, M. (2016). Morphology predicts species’ functional roles and their degree of specialization in plant–Frugivore interactions. Proc. R. Soc. B Biol. Sci., 283.

Dongmo, M.A.K., Bonebrake, T.C., Hanna, R. & Fomena, A. (2018). Seasonal Polyphenism in Bicyclus dorothea (Lepidoptera: Nymphalidae) Across Different Habitats in Cameroon. Environ. Entomol., 47, 1601–1608.

Dormann, C.F., Frund, J., Bluthgen, N. & Gruber, B. (2009). Indices, Graphs and Null Models: Analyzing Bipartite Ecological Networks. Open Ecol. J., 2, 7–24.

Dupont, Y.L., Padrón, B., Olesen, J.M. & Petanidou, T. (2009). Spatio-temporal variation in the structure of pollination networks. Oikos, 118, 1261–1269.

Dupont, Y.L., Trøjelsgaard, K., Hagen, M., Henriksen, M. V., Olesen, J.M., Pedersen, N.M.E., et al. (2014). Spatial structure of an individual-based plant-pollinator network. Oikos, 123, 1301–1310.

Egas, M., Sabelis, M.W. & Dieckmann, U. (2005). Evolution of specialization and ecological character displacement of herbivores along a gradient of plant quality. Evolution (N. Y)., 59, 507–520.

Eklöf, A., Jacob, U., Kopp, J., Bosch, J., Castro-Urgal, R., Chacoff, N.P., et al. (2013). The dimensionality of ecological networks. Ecol. Lett., 16, 577–583.

Fenberg, P.B., Self, A., Stewart, J.R., Wilson, R.J. & Brooks, S.J. (2016). Exploring the universal ecological responses to climate change in a univoltine butterfly. J. Anim. Ecol., 85, 739–748.

Fort, H., Vázquez, D.P. & Lan, B.L. (2016). Abundance and generalisation in mutualistic networks: solving the chicken-and-egg dilemma. Ecol. Lett., 19, 4–11.

Fric, Z.K. & Konvicka, M. (2002). Generations of the polyphenic butterfly Araschnia levana differ in body design. Evol. Ecol. Res. Evolutionary Ecology, Ltd.

Fries, T.C.E. (1925). The vertical distribution of some plants on Nuolja (Torne Lappmark). Bot. Not., 1925, 205–216.

Fukami, T., Martijn Bezemer, T., Mortimer, S.R. & van der Putten, W.H. (2005). Species divergence and trait convergence in experimental plant community assembly. Ecol. Lett., 8, 1283–1290.

Gentile, G., Bonelli, S. & Riva, F. (2021). Evaluating intraspecific variation in insect trait analysis. Ecol. Entomol., 46, 11–18.

Germain, R.M., Williams, J.L., Schluter, D. & Angert, A.L. (2018). Moving Character Displacement beyond Characters Using Contemporary Coexistence Theory. Trends Ecol. Evol., 33, 74–84.

Gillespie, M.A.K., Birkemoe, T. & Sverdrup-Thygeson, A. (2017). Interactions between body size, abundance, seasonality, and phenology in forest beetles. Ecol. Evol., 7, 1091–1100.

Gómez, J.M., Perfectti, F., Armas, C., Narbona, E., González-Megías, A., Navarro, L., et al. (2020). Within-individual phenotypic plasticity in flowers fosters pollination niche shift. Nat. Commun., 11, 1–12.

Gravel, D., Baiser, B., Dunne, J.A., Kopelke, J.-P., Martinez, N.D., Nyman, T., et al. (2019). Bringing Elton and Grinnell together: a quantitative framework to represent the biogeography of ecological interaction networks. Ecography (Cop.)., 42, 401–415.

Gray, R.E.J., Ewers, R.M., Boyle, M.J.W., Chung, A.Y.C. & Gill, R.J. (2018). Effect of tropical forest disturbance on the competitive interactions within a diverse ant community. Sci. Rep., 8, 1–12.

Greenleaf, S.S., Williams, N.M., Winfree, R. & Kremen, C. (2007). Bee foraging ranges and their relationship to body size. Oecologia, 153, 589–596.

Günter, F., Beaulieu, M., Brunetti, M., Lange, L., Schmitz Ornés, A. & Fischer, K. (2019). Latitudinal and altitudinal variation in ecologically important traits in a widespread butterfly. Biol. J. Linn. Soc., 128, 742–755.

Guthrie, RD. (2003). Rapid body size decline in Alaskan Pleistocene horses before extinction. Nat., 426, 169–171.

Gutiérrez, D. & Wilson, R.J. (2021). Intra-and interspecific variation in the responses of insect phenology to climate. J. Anim. Ecol., 90, 248–259.

Hagen, M., Kissling, W.D., Rasmussen, C., De Aguiar, M.A.M., Brown, L.E., Carstensen, D.W., et al. (2012). Biodiversity, Species Interactions and Ecological Networks in a Fragmented World. In: Advances in Ecological Research. Academic Press Inc., pp. 89–210.

Heinrich, B. (1975). Thermoregulation in Bumblebees II. Energetics of Warm-up and Free Flight. J. comp. Physiol. Springer-Verlag.

Horne, C.R., Hirst, A.G. & Atkinson, D. (2015). Temperature-size responses match latitudinal-size clines in arthropods, revealing critical differences between aquatic and terrestrial species. Ecol. Lett., 18, 327–335.

Hurtado, P., Prieto, M., Martínez-Vilalta, J., Giordani, P., Aragón, G., López-Angulo, J., et al. (2020). Disentangling functional trait variation and covariation in epiphytic lichens along a continent-wide latitudinal gradient. Proc. R. Soc. B Biol. Sci., 287.

Ings, T.C., Montoya, J.M., Bascompte, J., Blüthgen, N., Brown, L., Dormann, C.F., et al. (2009). Review: Ecological networks - beyond food webs. J. Anim. Ecol., 78, 253–269.

Jourdan, J., Krause, S.T., Lazar, V.M., Zimmer, C., Sommer-Trembo, C., Arias-Rodriguez, L., et al. (2016). Shared and unique patterns of phenotypic diversification along a stream gradient in two congeneric species. Sci. Rep., 6, 1–20.

Kaiser-Bunbury, C.N., Valentin, T., Mougal, J., Matatiken, D. & Ghazoul, J. (2011). The tolerance of island plant-pollinator networks to alien plants. J. Ecol., 99, 202–213.

Kenna, D., Pawar, S. & Gill, R.J. (2021). Thermal flight performance reveals impact of warming on bumblebee foraging potential. Funct. Ecol.

Keyes, A.A., McLaughlin, J.P., Barner, A.K. & Dee, L.E. (2021). An ecological network approach to predict ecosystem service vulnerability to species losses. Nat. Commun., 12, 1–11.

Klumpers, S.G.T., Stang, M. & Klinkhamer, P.G.L. (2019). Foraging efficiency and size matching in a plant–pollinator community: the importance of sugar content and tongue length. Ecol. Lett., 22, 469–479.

Kortsch, S., Primicerio, R., Fossheim, M., Dolgov, A. V. & Aschan, M. (2015). Climate change alters the structure of arctic marine food webs due to poleward shifts of boreal generalists. Proc. R. Soc. B Biol. Sci., 282.

Kuppler, J., Höfers, M.K., Wiesmann, L. & Junker, R.R. (2016). Time-invariant differences between plant individuals in interactions with arthropods correlate with intraspecific variation in plant phenology, morphology and floral scent. New Phytol., 210, 1357–1368.

Larsen, T.H., Williams, N.M. & Kremen, C. (2005). Extinction order and altered community structure rapidly disrupt ecosystem functioning. Ecol. Lett., 8, 538–547.

Lecoq, L., Ernoult, A. & Mony, C. (2021). Past landscape structure drives the functional assemblages of plants and birds. Sci. Rep., 11, 3443.

Lundberg, H. & Ranta, E. (1980). Habitat and Food Utilization in a Subarctic Bumblebee Community. Oikos, 35, 303.

MacDougall, A.S., Caplat, P., Olofsson, J., Siewert, M.B., Bonner, C., Esch, E., et al. (2021). Comparison of the distribution and phenology of Arctic Mountain plants between the early 20th and 21st centuries. Glob. Chang. Biol., 00, 1–14.

Maechler, M., Rousseeuw, P., Struyf, A., Hubert, M., Hornik, K., Studer, M., et al. (2014). Package “cluster.”

McCabe, L.M., Cobb, N.S. & Butterfield, B.J. (2019). Environmental filtering of body size and darker coloration in pollinator communities indicate thermal restrictions on bees, but not flies, at high elevations. PeerJ, 2019, e7867.

McGill, B.J., Enquist, B.J., Weiher, E. & Westoby, M. (2006). Rebuilding community ecology from functional traits. Trends Ecol. Evol., 21, 178–185.

Meredith, M. & Ridout, M. (2020). Overview of the overlap package.

Miller-Struttmann, N.E. & Galen, C. (2014). High-altitude multi-taskers: bumble bee food plant use broadens along an altitudinal productivity gradient. Oecologia, 1, 1033–1045.

Miller-Struttmann, N.E., Geib, J.C., Franklin, J.D., Kevan, P.G., Holdo, R.M., Ebert-May, D., et al. (2015). Functional mismatch in a bumble bee pollination mutualism under climate change. Science (80-.)., 349, 1541–1544.

Ohler, L.-M., Lechleitner, M. & Junker, R.R. (2020). Microclimatic effects on alpine plant communities and flower-visitor interactions. Sci. Rep., 10, 1–9.

Oksanen, J., Kindt, R., Legendre, P., O’Hara, B., Stevens, M.H.H., Oksanen, M.J., et al. (2007). The vegan package. Community Ecol. Packag., 10, 719.

Olesen, J.M., Bascompte, J., Elberling, H. & Jordano, P. (2008). TEMPORAL DYNAMICS IN A POLLINATION NETWORK. Ecology, 89, 1573–1582.

Orr, M.C., Hughes, A.C., Chesters, D., Pickering, J., Zhu, C.D. & Ascher, J.S. (2021). Global Patterns and Drivers of Bee Distribution. Curr. Biol., 31, 451-458.e4.

Pellissier, L., Albouy, C., Bascompte, J., Farwig, N., Graham, C., Loreau, M., et al. (2018). Comparing species interaction networks along environmental gradients. Biol. Rev., 93, 785–800.

Peralta, G., Vázquez, D.P., Chacoff, N.P., Lomáscolo, S.B., Perry, G.L.W. & Tylianakis, J.M. (2020). Trait matching and phenological overlap increase the spatio-temporal stability and functionality of plant–pollinator interactions. Ecol. Lett., 23, 1107–1116.

Pigot, A.L., Sheard, C., Miller, E.T., Bregman, T.P., Freeman, B.G., Roll, U., et al. (2020). Macroevolutionary convergence connects morphological form to ecological function in birds. Nat. Ecol. Evol., 4, 230–239.

Pinheiro, J., Bates, D., DebRoy, S., Sarkar, D., Heisterkamp, S., Van Willigen, B., et al. (2017). Package ‘nlme.’ Linear nonlinear Mix. Eff. Model. version, 3.

Poisot, T., Stouffer, D.B. & Kéfi, S. (2016). Describe, understand and predict: why do we need networks in ecology?

R Core Team (2020). R: A language and environment for statistical computing. R Foundation for Statistical Computing, Vienna, Austria. URL https://www.R-project.org/.

Ranta, E. & Lundberg, H. (1981). Resource utilization by bumblebee queens, workers and males in a subarctic area. Ecography (Cop.)., 4, 145–154.

Ridout, M.S. & Linkie, M. (2009). Estimating overlap of daily activity patterns from camera trap data. J. Agric. Biol. Environ. Stat., 14, 322–337.

Robroek, B.J.M., Jassey, V.E.J., Payne, R.J., Martí, M., Bragazza, L., Bleeker, A., et al. (2017). Taxonomic and functional turnover are decoupled in European peat bogs. Nat. Commun., 8, 1–9.

Des Roches, S., Post, D.M., Turley, N.E., Bailey, J.K., Hendry, A.P., Kinnison, M.T., et al. (2018). The ecological importance of intraspecific variation. Nat. Ecol. Evol., 2, 57–64.

Rode, K.D., Amstrup, S.C. & Regehr, E. V. (2010). Reduced body size and cub recruitment in polar bears associated with sea ice decline. Ecol. Appl., 20, 768–782.

Rohlf, F.J. (2015). The tps series of software. Hystrix, 26, 1–4.

Rumeu, B., Sheath, D.J., Hawes, J.E. & Ings, T.C. (2018). Zooming into plant-flower visitor networks: An individual trait-based approach. PeerJ, 2018, e5618.

Schroeder, P.J. & Jenkins, D.G. (2018). How robust are popular beta diversity indices to sampling error? Ecosphere, 9, e02100.

Siefert, A., Violle, C., Chalmandrier, L., Albert, C.H., Taudiere, A., Fajardo, A., et al. (2015). A global meta-analysis of the relative extent of intraspecific trait variation in plant communities. Ecol. Lett., 18, 1406–1419.

Signorell, A. (2021). DescTools: Tools for Descriptive Statistics.

Söderström, B. (2017). Sveriges humlor: en fälthandbok. Entomologiska föreningen i Stockholm.

Soroye, P., Newbold, T. & Kerr, J. (2020). Climate change contributes to widespread declines among bumble bees across continents. Science (80)., 367, 685–688.

Stang, M., Klinkhamer, P.G.L., Waser, N.M., Stang, I. & van der Meijden, E. (2009). Size-specific interaction patterns and size matching in a plant–pollinator interaction web. Ann. Bot., 103, 1459–1469.

Taylor-Cox, E.D., Macgregor, C.J., Corthine, A., Hill, J.K., Hodgson, J.A. & Saccheri, I.J. (2020). Wing morphological responses to latitude and colonisation in a range expanding butterfly. PeerJ, 8, e10352.

Thuiller, W., Pironon, S., Psomas, A., Barbet-Massin, M., Jiguet, F., Lavergne, S., et al. (2014). The European functional tree of bird life in the face of global change. Nat. Commun., 5, 1–10.

Trøjelsgaard, K. & Olesen, J.M. (2016). Ecological networks in motion: micro-and macroscopic variability across scales. Funct. Ecol.

Tur, C., Vigalondo, B., Trøjelsgaard, K., Olesen, J.M. & Traveset, A. (2014). Downscaling pollen–transport networks to the level of individuals. J. Anim. Ecol., 83, 306–317.

Tylianakis, J.M. & Morris, R.J. (2017). Ecological Networks Across Environmental Gradients. Annu. Rev. Ecol. Evol. Syst., 48, 25–48.

Vázquez, D.P., Melián, C.J., Williams, N.M., Blüthgen, N., Krasnov, B.R. & Poulin, R. (2007). Species abundance and asymmetric interaction strength in ecological networks. Oikos, 116, 1120–1127.

Wardle, D.A. & Zackrisson, O. (2005). Effects of species and functional group loss on island ecosystem properties. Nat. 2005 4357043, 435, 806–810.

Williams, P.H., Byvaltsev, A.M., Cederberg, B., Berezin, M. V., Ødegaard, F., Rasmussen, C., et al. (2015). Genes suggest ancestral colour polymorphisms are shared across morphologically cryptic species in arctic bumblebees. PLoS One, 10, 1–26.

Wong, M.K.L. & Carmona, C.P. (2021). Including intraspecific trait variability to avoid distortion of functional diversity and ecological inference: Lessons from natural assemblages. Methods Ecol. Evol., 12, 946–957.

Wu, C.-H., Holloway, J.D., Hill, J.K., Thomas, C.D., Chen, I.-C. & Ho, C.-K. (2019). Reduced body sizes in climate-impacted Borneo moth assemblages are primarily explained by range shifts. Nat. Commun. 2019 101, 10, 1–7.

