## Supplementary Information for "Mapping trait versus species turnover reveals spatiotemporal variation in functional redundancy in a plant-pollinator network"

Cantwell-Jones *et al.*

**SUPPLEMENTARY INFORMATION**

**Supplementary Methods**

Supplementary Information 1: **Sampling plots details and set-up**

The 3.4 km transect was divided into 79 posts located approximately 45.0 m apart (range: 34.4-59.0 m) and spanned five vegetation zones. Plots used for surveying were established in 2018, with geographical coordinates and altitude logged with a high-precision GPS (Trimble dGPS T10 tables and R8s GNSS) and marked in field with flags and marker tape. A total of 13 plots were spread across the transect and altitudinal gradient, covering the five vegetation zones. Initial assessment of Mount Nuolja discounted plots considered unsuitable for surveying (e.g. plots positioned on a cliff edge). The remaining plots were randomly selected ensuring an equal representation of each vegetation zone (≥2 plots per vegetation zone).

Each plots was a 45 × 45 m quadrat established parallel to the transect, with the orientation of the plot (left or right of the transect) chosen randomly. The plots were positioned 6 m away from the main transect line to minimise the effects of human interference (such as, trampled vegetation).

Supplementary Information 2: **Bumblebee observation and sampling protocol**

We avoided sampling a given plot at the same time each day but randomly selecting which plot we start sampling on. On a given day, after sampling the first plot, we would continue sampling up the mountain, alternating the plots visited. Once the highest elevation plot was reached, we would return to the first sampled plot and sample plots down-elevation of it, again alternating the plots visited. For example, imagine the sampling plots were numbered one to 13, with 13 representing the highest elevation plot. If plot 9 was randomly selected to be the starting plot, we would sample plot 9 first, then plot 11 and 13. Then, we would sample plot 7, 5, 3 and 1. The remaining plots would be sampled the next day, again with the first plot being randomly selected. Any periods of rain or snow (≥0.2 mm hr^-1^) suspended field trips until the next suitable day.

Bumblebee surveys were conducted by three trained individuals. During the 20-min observational period, two individuals walked along the established bumblebee quadrat and caught bees, while the third person acted as a scribe. Scribes monitored surveys using a stopwatch, which was paused when bees were being transferred to holding pots, then to insulated bags. Bias in surveying ability was avoided by rotating the roles of surveyors and scribe between plots (Caro et al., 1979). Although observations within a plot were recorded for 20 min, the total amount of time spent surveying ranged from 20 min to 1 hour and 28 min, with a median duration of 26 min, due to the varying amount of time taken to transfer bumblebees to pots. After survey completion, bumblebees were transferred from the pots to a marking cage, where their species identity was confirmed and a digital photo taken (Canon SX720 HS). After this processing, bees were released.

Supplementary Information 3: **Assigning female castes**

All bees for which intertegular distance (ITD) was measured and caste was confidently assigned in the field (*n* = 138; 49 queens & 89 workers) were used to inform a model predicting female caste from ITD measures. Due to clear morphological differences, bees identified as males in the field were assumed correctly identified and bees of unknown caste (*n* = 773) assumed female. A linear mixed-effects model (lme; using package “lme4”; Bates et al., 2018) was used to identify differences in mean ITD between castes, while accounting for the presence of different bumblebee species (the random intercepts). Binomial logistic regression (glm) was then used to predict caste of the unknown bees using ITD based upon 95 % likelihood. Any female individuals belonging to species in the subgenus *Psithyrus* were assumed to be queens, as these considered parasitic species do not have a worker caste (Martin et al., 2010) and hence these individuals were excluded from the mixed-effects model and logistic regression. Based upon the likelihood predictions of this logistic regression, a further 573 bees could be assigned to a female caste based with greater than 95 % probability.

Supplementary Information 4: **Determining the cut-off for the “early” and “late” seasons.**

Within each spatially pooled set of observations, we split observations into an “early” and “late” part of the season. To guide this split we considered colony-lifecycle phenology of the social bumblebees (queens emerge and dominate during the early season and workers during the late) to be natural stages of comparison to look at temporal functional turnover. To determine the switch from foraging queens to foraging workers, we used a binomial generalised linear model (glm) with “logit” link, where the proportion of queens observed (out of all observed females) was regressed against the days since the start of sampling (df = 29; *z* = -2.29; *p* = 0.044; Table S7; Fig. S4). Subsequently, we determined the cut-off between the “early” and “late” part of the season as occurring when 50 % of observed female bees were likely to be queens (23^rd^ June 2018).

We intended this cut-off to be a means of separating the sampling season into two, and consequently we chose to use one date for the whole community (based on all the queens observed in the community), rather than creating a different date for each bumblebee species. Although species-specific dates would have allowed us to account for species differences in emergence times (and thus more accurate cut-offs for when a given species has more queens or workers), ten of the 13 species present in the community had ≤10 queens, making the fitting of a binomial generalised linear model challenging.

**Supplementary Methods Figures**

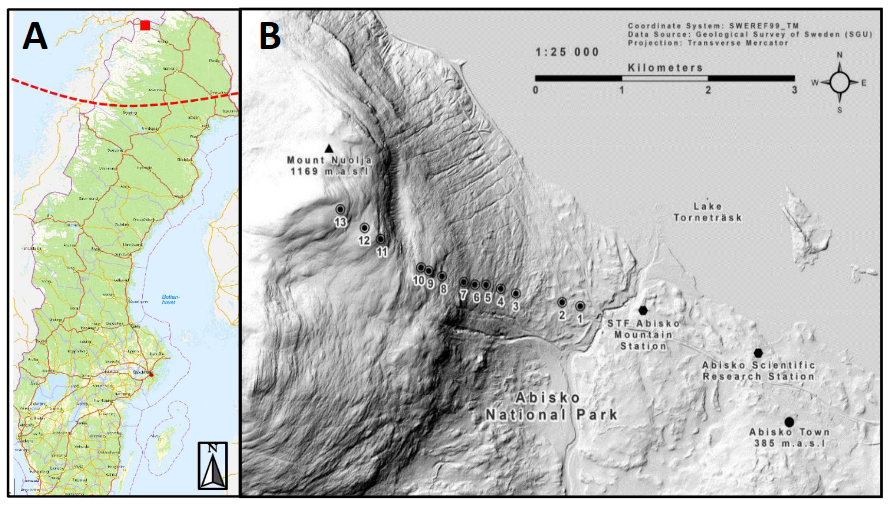

Supplementary Figure 1: **The location of the sampling transect A) in Sweden and B) on Mount Nuolja, Abisko National Park, showing the 13 sampling plots (numbered dots).** In A) the dashed red line indicates the Arctic circle. Figure taken from Cox (2018).

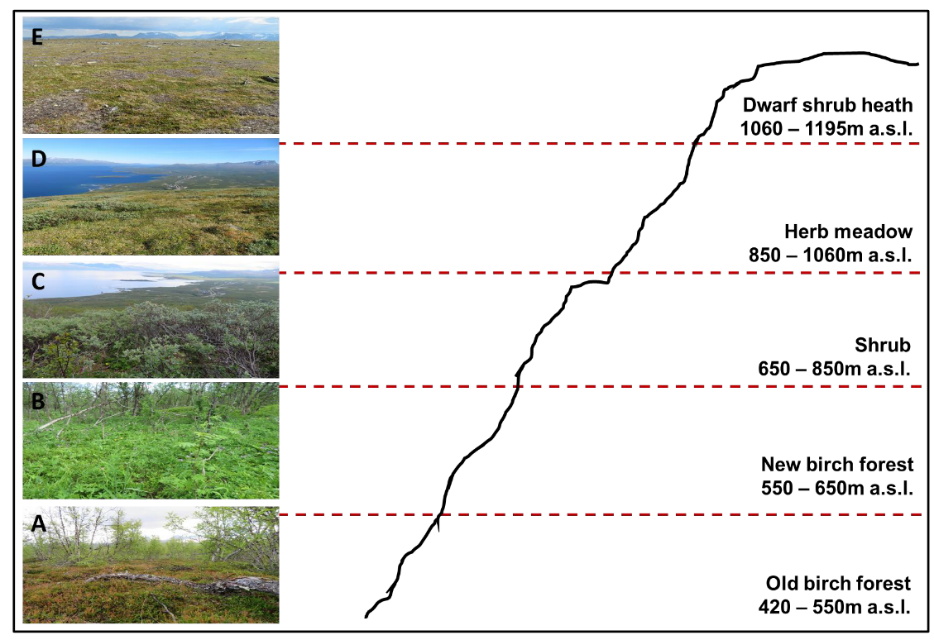

Supplementary Figure 2: **The five main vegetation types on Mount Nuolja.** Figure taken from Cox (2018). Vegetation zones A and B were classed as “low elevation”, and zones C, D and E, as “high elevation”.

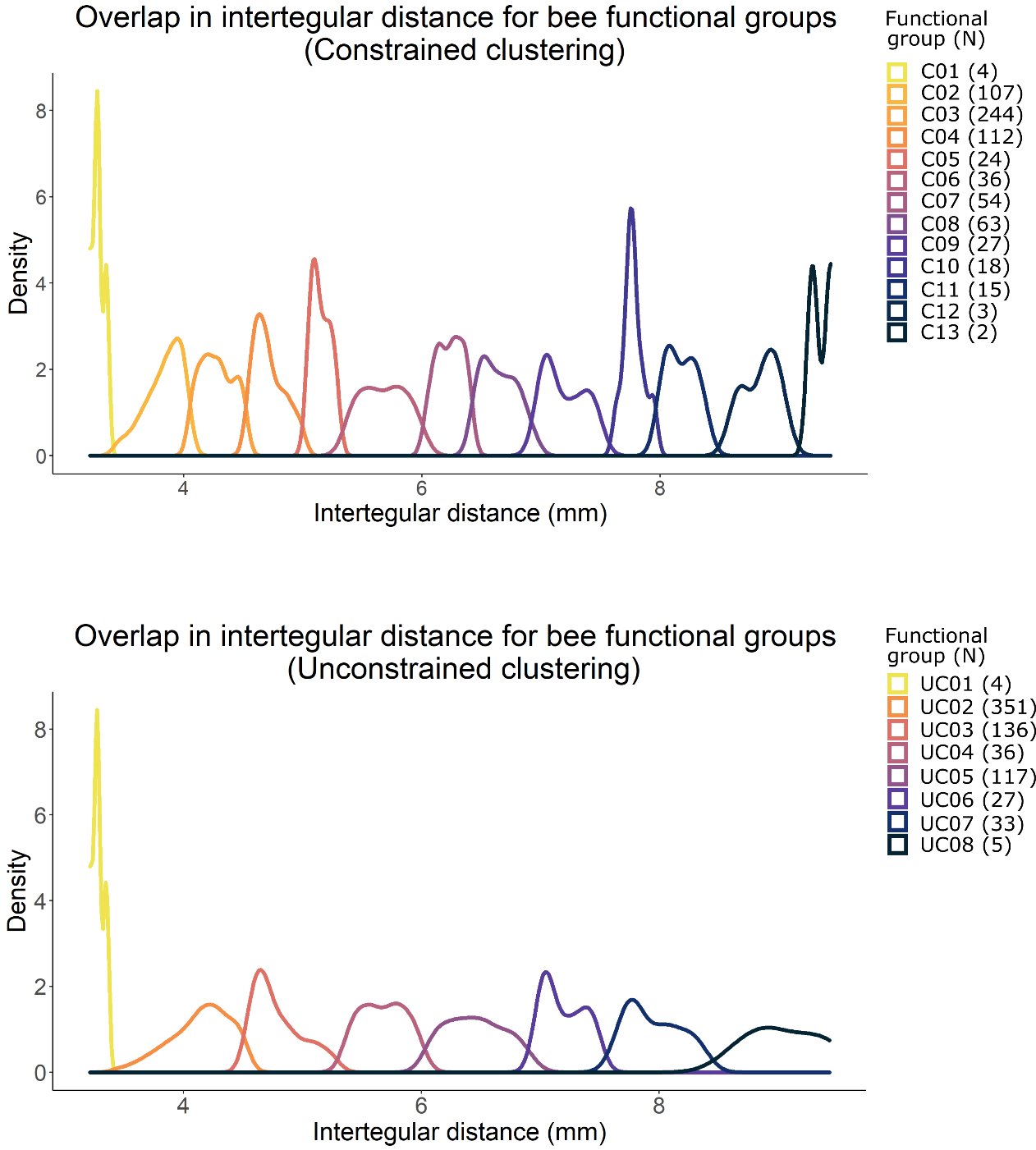

Supplementary Figure 3: **Overlap in intertegular distance for different bumblebee size functional groups when a constrained clustering approach was used.** Bumblebees were grouped into functional groups based on similarity in intertegular size and ignoring species identity. The sample size of each functional group is indicated in the legend.

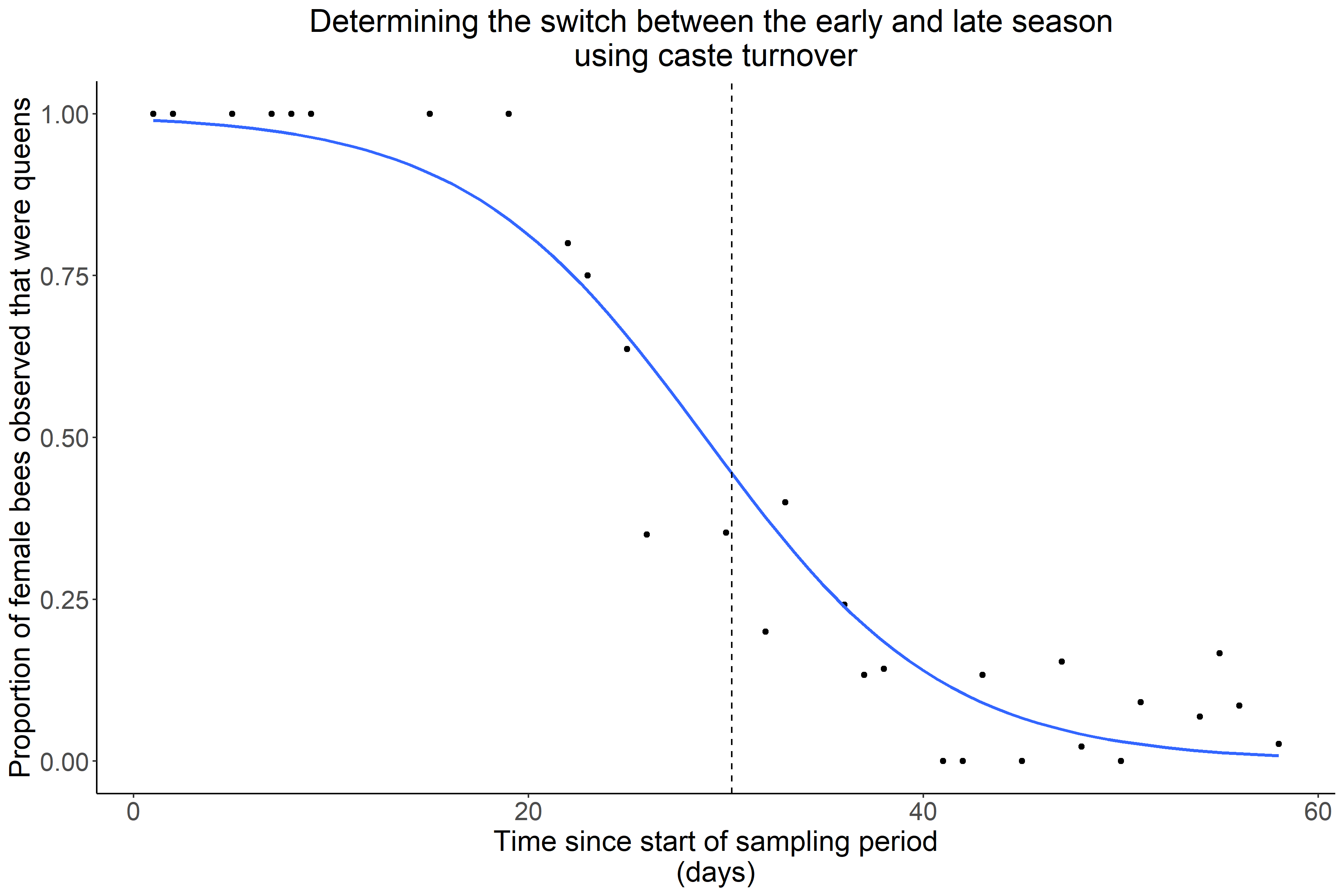

Supplementary Figure 4: **Determining the cut-off point for the “early” and “late” seasons using bumblebee lifecycle switch from queens foraging to workers foraging.** The date when 50 % of the female bees observed were queens (dashed line) was estimated using a binomial generalised linear model (solid blue line). This date (23/06/2018) was used as the cut-off point for the early and late seasons. *N* = 591 females.

**Supplementary Methods Tables**

| Supplementary Table 1: **The thirteen permanent sampling plots on Mount Nuolja.** The thirteen plots are denoted by the posts between which the plot lies. Total number of repeats = 217. Table taken from Cox (2018). | | | | | | | |
| --- | --- | --- | --- | --- | --- | --- | --- |
| Plot | Vegetation | Downslope compass bearing (⁰) | Upslope compass bearing (⁰) | Downslope elevation (m a.s.l.) | Upslope elevation (m a.s.l.) | Number of repeats | Earliest sampling date |
| 4-5 | Old birch (A) | 266 | 86 | 432 | 436 | 17 | 24/05/18 |
| 9-10 | Old birch (A) | 276 | 16 | 457 | 463 | 17 | 25/05/18 |
| 21-22 | Old birch (A) | 287 | 107 | 539 | 541 | 17 | 28/05/18 |
| 25-26 | New birch (B) | 282 | 102 | 578 | 587 | 17 | 25/05/18 |
| 29-30 | New birch (B) | 270 | 90 | 641 | 655 | 17 | 28/05/18 |
| 32-33 | Shrub (C) | 274 | 94 | 681 | 700 | 17 | 25/05/18 |
| 35-36 | Shrub (C) | 280 | 100 | 740 | 757 | 17 | 24/05/18 |
| 41-42 | Shrub (C) | 276 | 96 | 834 | 846 | 17 | 25/05/18 |
| 45-46 | Herb meadow (D) | 290 | 110 | 897 | 905 | 17 | 24/05/18 |
| 47-48 | Herb meadow (D) | 232 | 102 | 918 | 925 | 16 | 25/05/18 |
| 60-61 | Dwarf shrub heath (E) | 296 | 116 | 1,063 | 1,077 | 16 | 28/05/18 |
| 65-66 | Dwarf shrub heath (E) | 300 | 120 | 1,086 | 1,088 | 16 | 31/05/18 |
| 73-74 | Dwarf shrub heath (E) | 300 | 120 | 1,157 | 1,161 | 16 | 28/05/18 |

| Supplementary Table 2: **Summary of bee and flower species richness over space and time.** The bees included here were observed foraging and had their intertegular distance measured. | | | | |
| --- | --- | --- | --- | --- |
|  | Bee species richness | Flower species richness | Number of interactions | Constrained functional group richness |
| Early season, low elevation | 12 | 6 | 88 | 9 |
| Late season, low elevation | 12 | 16 | 224 | 10 |
| Early season, high elevation | 8 | 6 | 32 | 8 |
| Late season, high elevation | 11 | 21 | 151 | 12 |

| Supplementary Table 3: **Linear mixed-effects model testing for significant difference in intertegular distance (mm) between the queens and worker castes.** Bumblebee species were used as random intercepts. *N* = 138. Values in parentheses are the standard errors of the model estimates. Values in bold were considered statistically significant. Marginal R^2^ = 0.712; conditional R^2^ = 0.762. | | | |
| --- | --- | --- | --- |
| Coefficient | Estimate | *t*-value | *p*-value |
| Intercept (Queen caste) | 6.74  (± 0.131) | 51.4 | **<0.001** |
| Worker caste | -2.22  (± 0.118) | -18.8 | **<0.001** |
| Random effects | Variance | Standard deviation |  |
| *Bombus* species | 0.0812 | 0.285 |  |
| Residual | 0.381 | 0.617 |  |

| Supplementary Table 4: **Binomial generalized least squares model used for predicting the castes for female bees for whom the caste was unknown using intertegular distance.** *N* = 138. Values in parentheses are the standard errors of the model estimates. Values in bold were considered statistically significant. R^2^ = 0.828. | | | |
| --- | --- | --- | --- |
| Coefficient | Estimate | *z*-value | *p*-value |
| Intercept | 17.1  (± 2.83) | 6.04 | <0.001 |
| Intertegular distance  (mm) | -3.03  (±0.504) | -6.02 | <0.001 |

| Supplementary Table 5: **Summary of the data available for each *Bombus* species.** The sample size for each species is broken down into the different sexes and castes. “Unknown” caste refers to females who could not be assigned to the queen or worker caste with 95 % probability. The constrained and unconstrained functional groups refer to the different body-size-functional groups into which individuals were grouped, depending on whether constrained or unconstrained clustering was used, respectively. The data shown here refers to individuals that were observed foraging and had their intertegular distance (ITD) measured. | | | | | | |
| --- | --- | --- | --- | --- | --- | --- |
| Species | Sample size | | | | ITD (mm) | Constrained functional groups |
|  | Queens | Workers | Drones | Unknown |  |  |
| *B. alpinus/polaris* | 10 | 4 | 0 | 4 | 4.12-8.88 | C03, C04, C06, C07, C08, C09, C10, C11, C12 |
| *B. balteatus* | 9 | 6 | 0 | 14 | 3.74-8.34 | C02, C03, C04, C05, C06, C08, C09, C10, C11 |
| *B. bohemicus* | 2 | 0 | 0 | 2 | 5.10-7.65 | C05, C08, C09 |
| *B. cingulatus* | 3 | 1 | 0 | 0 | 3.93-6.58 | C02, C07, C08 |
| *B. flavidus* | 0 | 0 | 0 | 2 | 4.55-4.82 | C04 |
| *B. hortorum* | 3 | 10 | 0 | 6 | 3.59-7.77 | C02, C03, C04, C06, C07, C08, C09, C10 |
| *B. hyperboreus* | 1 | 0 | 0 | 0 | 7.47 | C09 |
| *B. jonellus* | 3 | 97 | 5 | 15 | 3.21-7.04 | C01, C02, C03, C04, C05, C06, C07, C08, C09 |
| *B. lapponicus* | 14 | 26 | 6 | 15 | 3.80-7.78 | C02, C03, C04, C05, C06, C07, C08, C09, C10 |
| *B. lucorum* | 13 | 8 | 0 | 10 | 4.27-9.28 | C03, C04, C05, C06, C07, C08, C09, C10, C11, C12, C13 |
| *B. monticola* | 8 | 24 | 4 | 14 | 3.60-7.46 | C02, C03, C04, C05, C07, C08, C09 |
| *B. pascuorum* | 5 | 5 | 0 | 3 | 3.92-7.01 | C02, C03, C04, C05, C06, C07, C08, C09 |
| *B. pratorum* | 13 | 89 | 6 | 23 | 3.27-8.67 | C01, C02, C03, C04, C06, C07, C08, C11, C12 |

| Supplementary Table 6: **Summary of the functional groups, assigned using constrained clustering.** The sample size for each functional group is broken down into the different sexes and castes. “Unknown” caste refers to females who could not be assigned to the queen or worker caste with 95 % probability. The data shown here refers to individuals that were observed foraging and had their intertegular distance (ITD) measured. Individuals across all points in space and time were pooled prior to clustering. | | | | | | |
| --- | --- | --- | --- | --- | --- | --- |
| Functional group | Sample size | | | | ITD (mm) | *Bombus* species |
|  | Queens | Workers | Drones | Unknown |  |  |
| C01 | 0 | 4 | 0 | 0 | 3.21-3.34 | *B. jonellus, B. pratorum* |
| C02 | 0 | 77 | 5 | 0 | 3.46-4.04 | *B. balteatus, B. cingulatus, B. hortorum, B. jonellus, B. lapponicus, B. monticola, B. pascuorum, B. pratorum* |
| C03 | 1 | 172 | 10 | 0 | 4.04-4.52 | *B. alpinus/polaris, B. balteatus, B. hortorum, B. jonellus, B. lapponicus, B. lucorum, B. monticola, B. pascuorum, B. pratorum* |
| C04 | 1 | 13 | 4 | 63 | 4.53-4.99 | *B. alpinus/polaris, B. balteatus, B. flavidus, B. hortorum, B. jonellus, B. lapponicus, B. lucorum, B. monticola, B. pascuorum, B. pratorum* |
| C05 | 0 | 2 | 1 | 14 | 5.03-5.30 | *B. balteatus, B. bohemicus, B. jonellus, B. lapponicus, B. lucorum, B. monticola, B. pascuorum* |
| C06 | 4 | 1 | 0 | 17 | 5.39-5.98 | *B. alpinus/polaris, B. balteatus, B. hortorum, B. jonellus, B. lapponicus, B. lucorum, B. pascuorum, B. pratorum* |
| C07 | 11 | 0 | 0 | 16 | 6.05-6.40 | *B. alpinus/polaris, B. cingulatus, B. hortorum, B. jonellus, B. lapponicus, B. lucorum, B. monticola, B. pascuorum, B. pratorum* |
| C08 | 36 | 1 | 0 | 0 | 6.46-6.97 | *B. alpinus/polaris, B. balteatus, B. bohemicus, B. cingulatus, B. hortorum, B. jonellus, B. lapponicus, B. lucorum, B. monticola, B. pascuorum, B. pratorum* |
| C09 | 15 | 0 | 0 | 0 | 7.01-7.47 | *B. alpinus/polaris, B. balteatus, B. bohemicus, B. hortorum, B. hyperboreus, B. jonellus, B. lapponicus, B. lucorum, B. monticola, B. pascuorum* |
| C10 | 11 | 0 | 0 | 0 | 7.71-7.95 | *B. alpinus/polaris, B. balteatus, B. hortorum, B. lapponicus, B. lucorum* |
| C11 | 11 | 0 | 1 | 0 | 8.02-8.34 | *B. alpinus/polaris, B. balteatus, B. lucorum, B. pratorum* |
| C12 | 3 | 0 | 0 | 0 | 8.67-8.99 | *B. alpinus/polaris, B. lucorum, B. pratorum* |
| C13 | 1 | 0 | 0 | 0 | 9.28 | *B. lucorum* |

| Supplementary Table 7: **Determining the cut-off point for the “early” and “late” seasons using bumblebee lifecycle switch from queens foraging to workers foraging.** A binomial generalised linear model (with “logit” link) was used to model the proportion of females that were queens over the course of the sampling period. Values in parentheses are the standard errors of the estimates; bolded values are statistically significant (*p* <0.05). *N* = 591 females. | | | |
| --- | --- | --- | --- |
| Coefficent | Estimate  (± standard error) | *z*-value | *p*-value |
| Intercept | 4.76  (± 1.86) | 2.55 | **0.0107** |
| Days since start of sampling | -1.16  (± 0.0577) | -2.29 | **0.044** |

**Supplementary Results Figures**

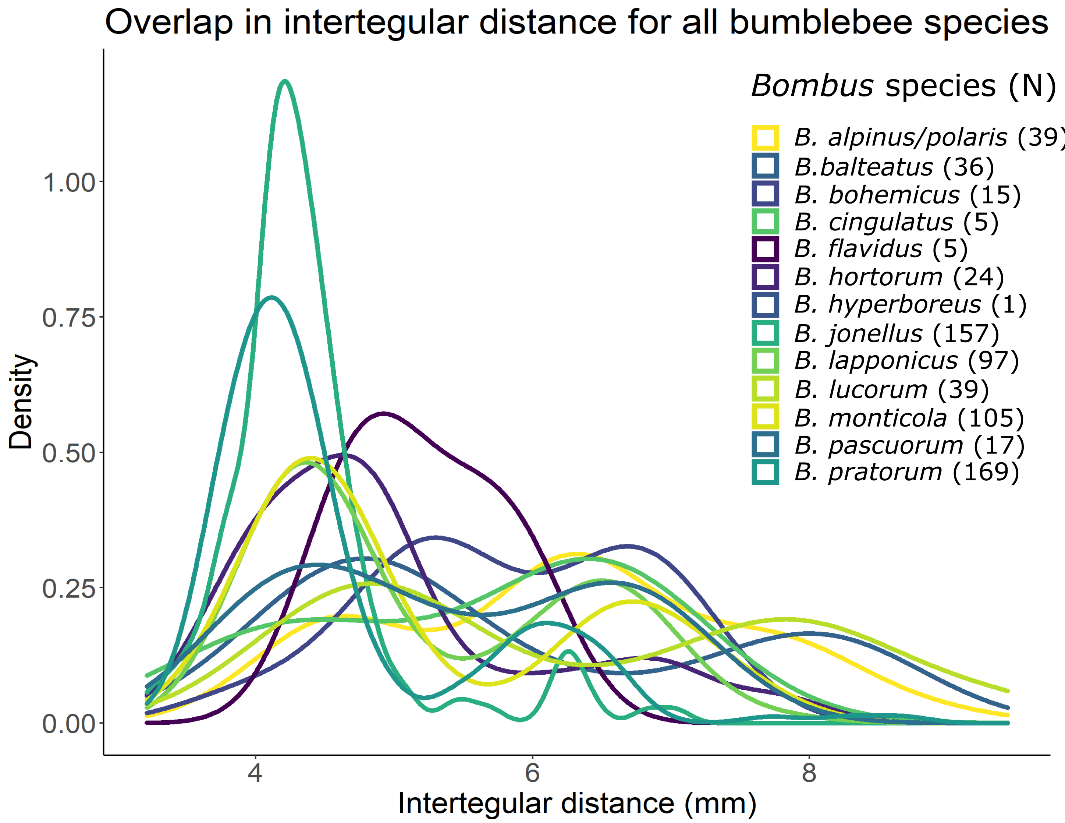

Supplementary Figure 5: **Overlap in intertegular distance (thorax width) across bumblebee species.** The sample size of each species is indicated in the legend.

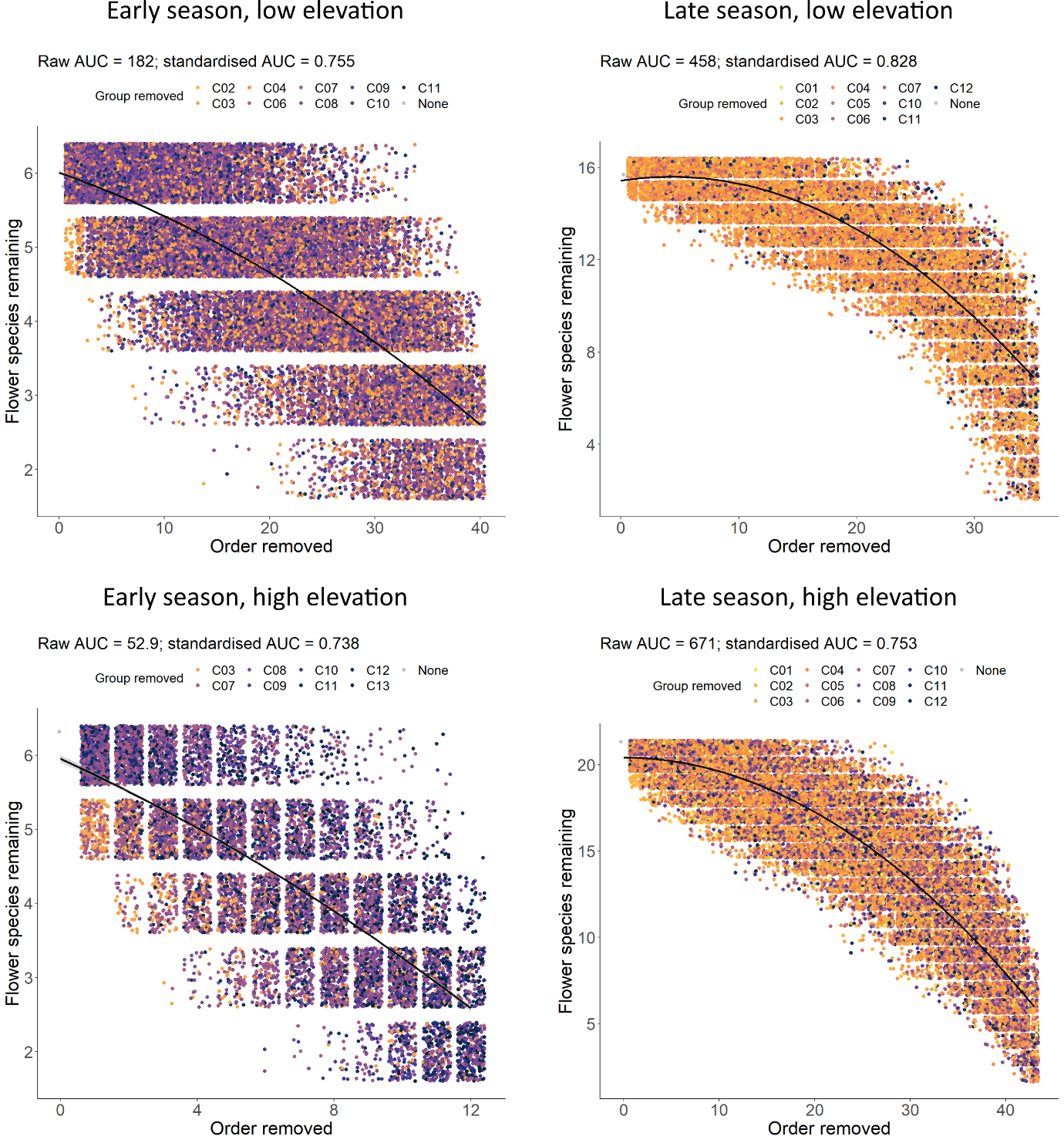

Supplementary Figure 6: **Change in number of remaining flower species in the network when constrained functional groups are randomly and successively removed from nodes (“Order removed”) on a plant-bumblebee-species network.** The random removal of functional groups until network collapse was iterated 1,000 times. Note that points on each plot were jittered. The standardised area under the curve (AUC) was calculated by estimating the AUC (based on the black regression line) and dividing this value by the product of the maximum number of flower species and functional groups removed.

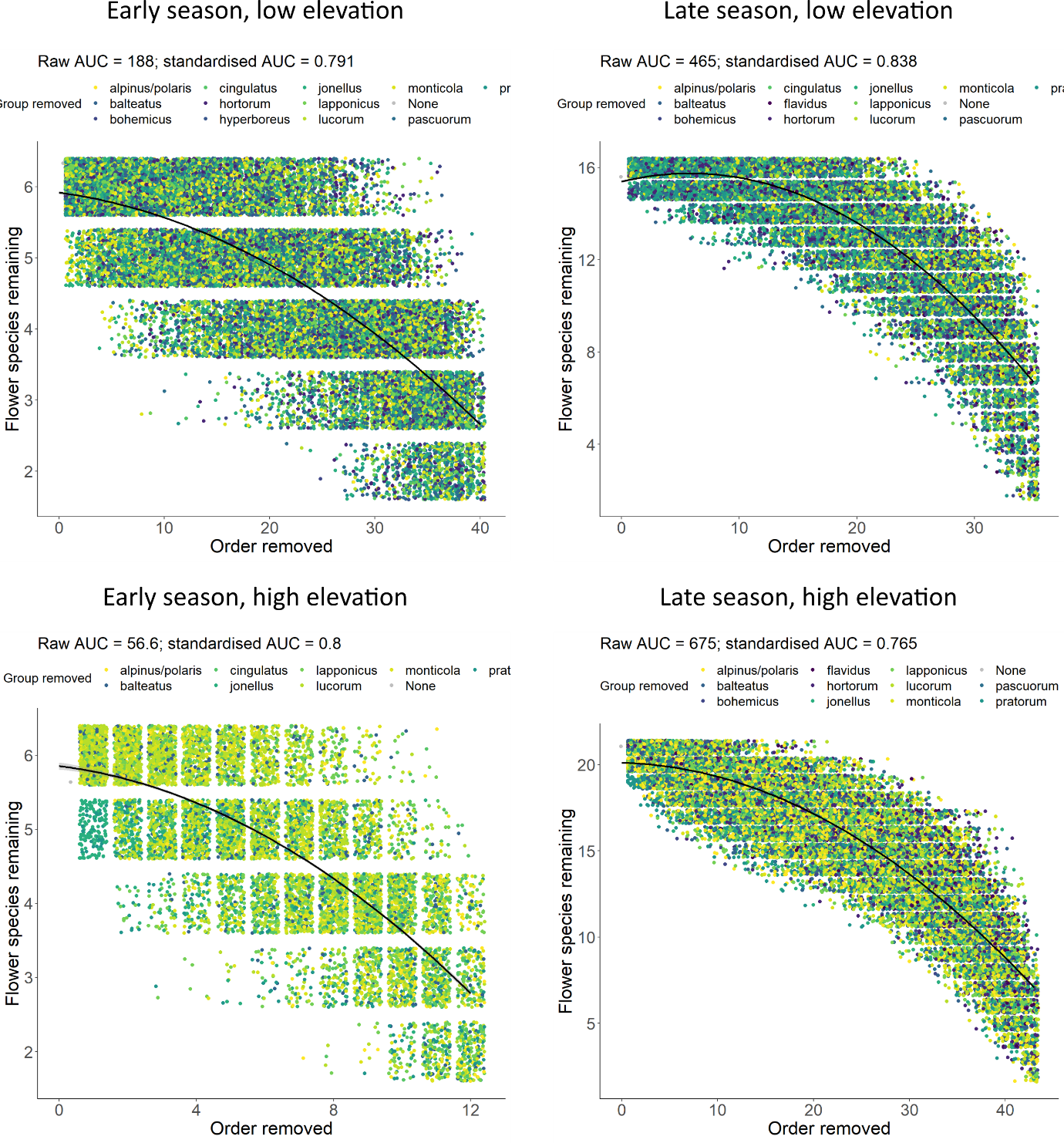

Supplementary Figure 7: **Change in number of remaining flower species in the network when bumblebee species are randomly and successively removed from constrained-functional-group nodes (“Order removed”) on a plant-bumblebee network.** The random removal of species until network collapse was iterated 1,000 times. Note that points on each plot were jittered. The standardised area under the curve (AUC) was calculated by estimating the AUC (based on the black regression line) and dividing this value by the product of the maximum number of flower species and species removed.

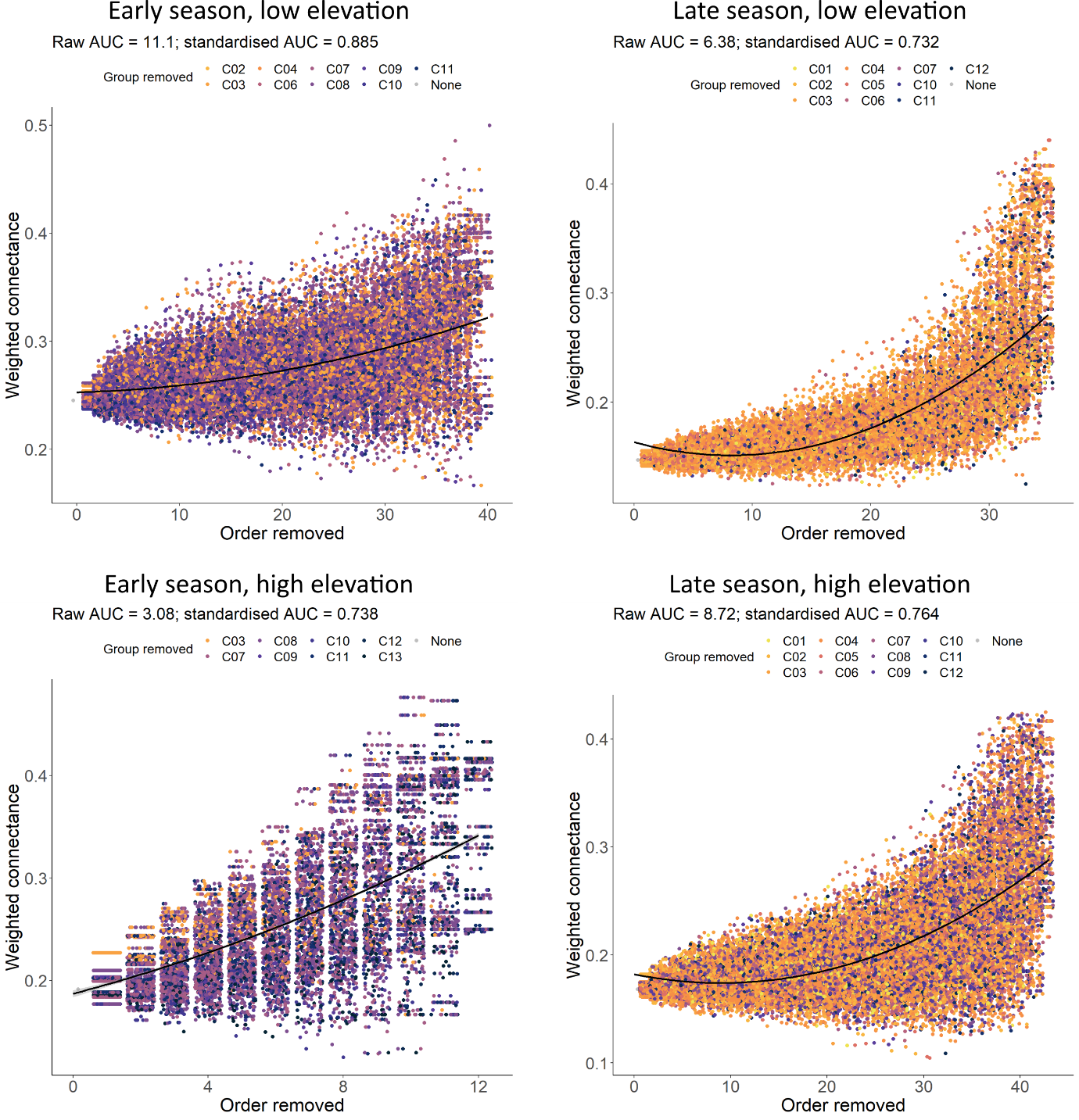

Supplementary Figure 8: **Change in the weighted connectance of each network when constrained functional groups are randomly and successively removed from nodes (“Order removed”) on a plant-bumblebee-species network.** The random removal of functional groups until network collapse was iterated 1,000 times. Note that points on each plot were jittered. The standardised area under the curve (AUC) was calculated by estimating the AUC (based on the black regression line) and dividing this value by the product of the maximum weighted connectance and functional groups removed.

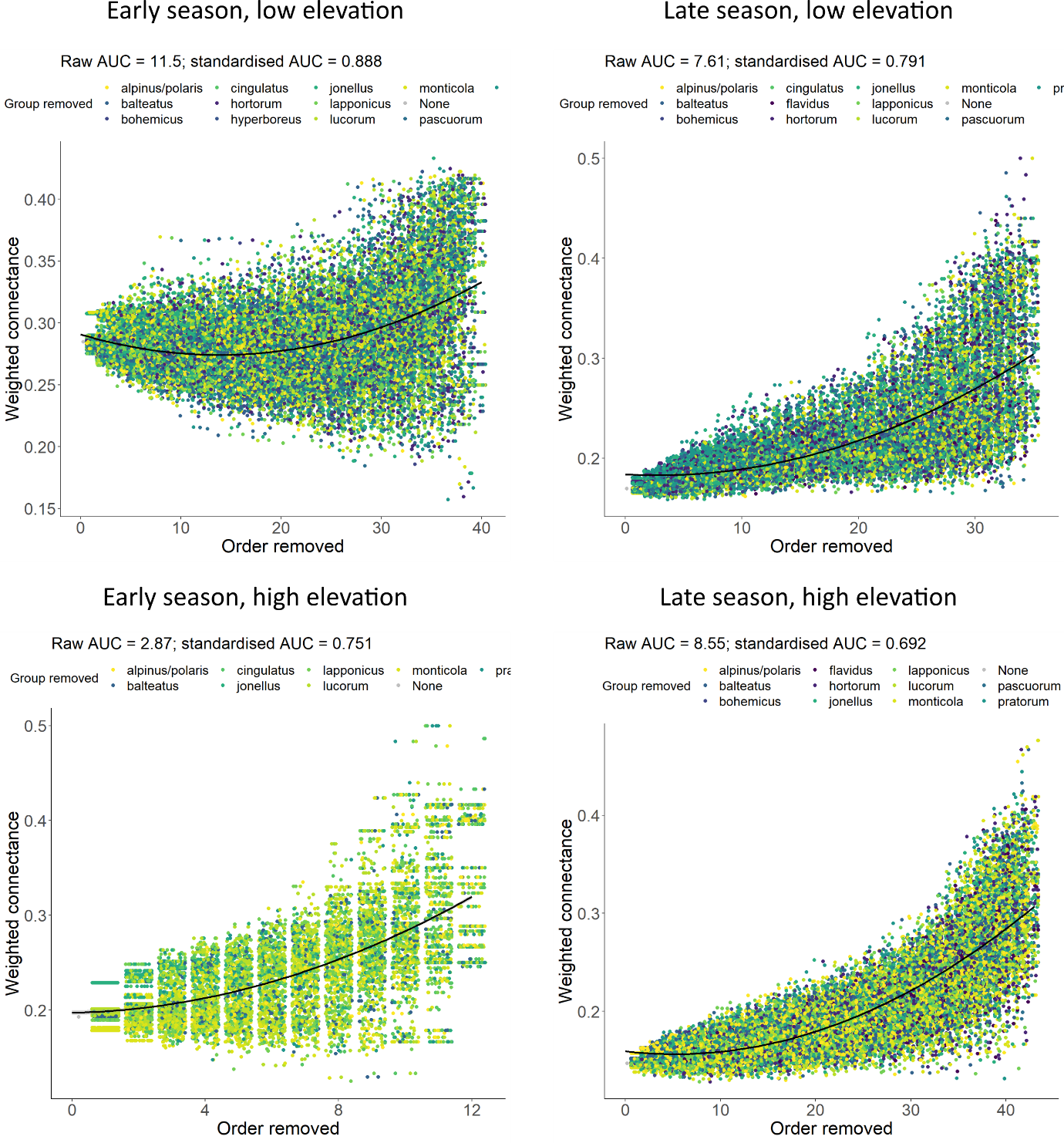

Supplementary Figure 9: **Change in the weighted connectance of the networks when bumblebee species are randomly and successively removed from constrained-functional-group nodes (“Order removed”) on a plant-bumblebee network.** The random removal of species until network collapse was iterated 1,000 times. Note that points on each plot were jittered. The standardised area under the curve (AUC) was calculated by estimating the AUC (based on the black regression line) and dividing this value by the product of the maximum weighted connectance and species removed.

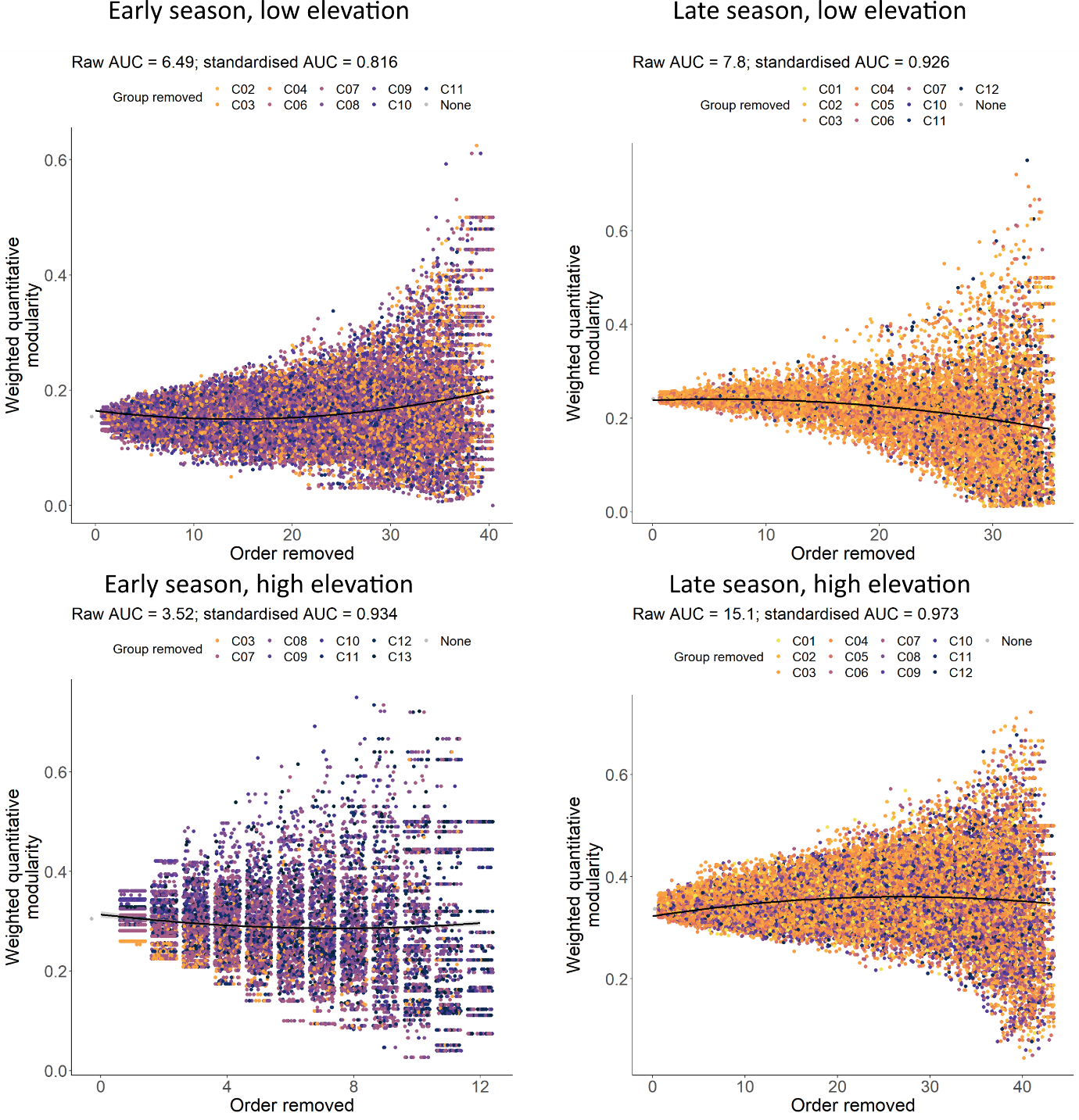

Supplementary Figure 10: **Change in the weighted quantitative modularity of each network when constrained functional groups are randomly and successively removed from nodes (“Order removed”) on a plant-bumblebee-species network.** The random removal of functional groups until network collapse was iterated 1,000 times. Note that points on each plot were jittered. The standardised area under the curve (AUC) was calculated by estimating the AUC (based on the black regression line) and dividing this value by the product of the maximum weighted quantitative modularity and functional groups removed.

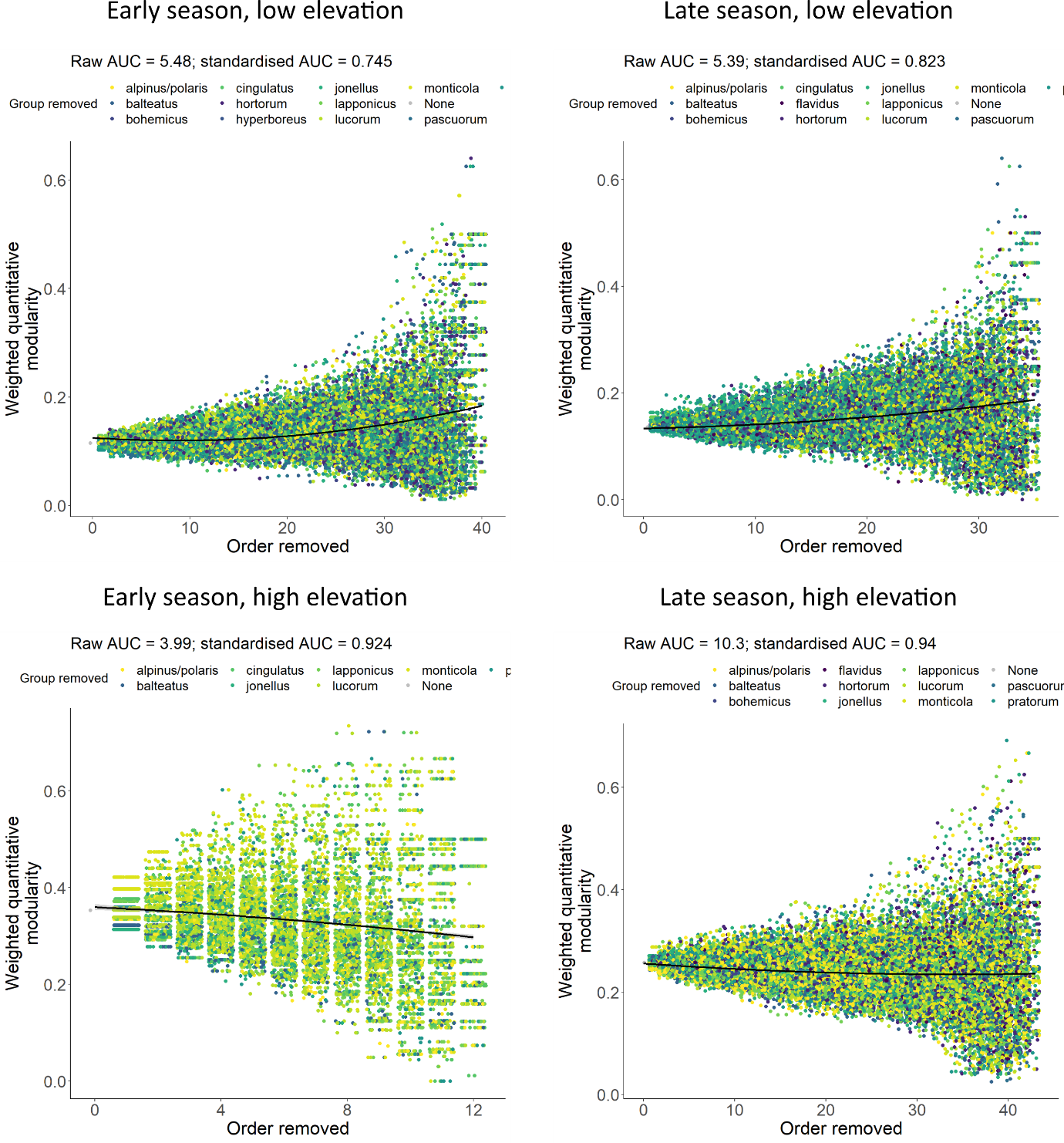

Supplementary Figure 11: **Change in the weighted quantitative modularity of the networks when bumblebee species are randomly and successively removed from constrained-functional-group nodes (“Order removed”) on a plant-bumblebee network.** The random removal of species until network collapse was iterated 1,000 times. Note that points on each plot were jittered. The standardised area under the curve (AUC) was calculated by estimating the AUC (based on the black regression line) and dividing this value by the product of the maximum weighted quantitative modularity and species removed.

**Supplementary Results Tables**

| Supplementary Table 8: **Mixed-effects ANOVA assessing the ability of bumblebee species identity to explain differences in intertegular distance.** Castes nested within bumblebee species were included as random intercepts. | | | | |
| --- | --- | --- | --- | --- |
| Predictor | Sum of squares | Mean squares | *F*-value | *P*-value |
| *Bombus* species | 1.12 | 0.0934 | 0.362 | 0.967 |

| Supplementary Table 9: **Quantifying the degree of overlap in pairwise bumblebee species intertegular distance (mm) using the coefficient of overlapping.** The coefficient of overlapping compares the kernel density estimates of the two species. Species were included if they had ≥10 measurements of intertegular distance, yielding 45 pairwise combinations. | | |
| --- | --- | --- |
| *Bombus* species 1 | *Bombus* species 2 | Overlap |
| *B. balteatus* | *B. alpinus/polaris* | 0.637 |
| *B. bohemicus* | *B. alpinus/polaris* | 0.575 |
| *B. bohemicus* | *B. balteatus* | 0.527 |
| *B. hortorum* | *B. alpinus/polaris* | 0.522 |
| *B. hortorum* | *B. balteatus* | 0.683 |
| *B. hortorum* | *B. bohemicus* | 0.492 |
| *B. jonellus* | *B. alpinus/polaris* | 0.328 |
| *B. jonellus* | *B. balteatus* | 0.416 |
| *B. jonellus* | *B. bohemicus* | 0.251 |
| *B. jonellus* | *B. hortorum* | 0.609 |
| *B. lapponicus* | *B. alpinus/polaris* | 0.655 |
| *B. lapponicus* | *B. balteatus* | 0.565 |
| *B. lapponicus* | *B. bohemicus* | 0.581 |
| *B. lapponicus* | *B. hortorum* | 0.741 |
| *B. lapponicus* | *B. jonellus* | 0.638 |
| *B. lucorum* | *B. alpinus/polaris* | 0.672 |
| *B. lucorum* | *B. balteatus* | 0.856 |
| *B. lucorum* | *B. bohemicus* | 0.507 |
| *B. lucorum* | *B. hortorum* | 0.609 |
| *B. lucorum* | *B. jonellus* | 0.340 |
| *B. lucorum* | *B. lapponicus* | 0.500 |
| *B. monticola* | *B. alpinus/polaris* | 0.599 |
| *B. monticola* | *B. balteatus* | 0.588 |
| *B. monticola* | *B. bohemicus* | 0.539 |
| *B. monticola* | *B. hortorum* | 0.753 |
| *B. monticola* | *B. jonellus* | 0.648 |
| *B. monticola* | *B. lapponicus* | 0.888 |
| *B. monticola* | *B. lucorum* | 0.537 |
| *B. pascuorum* | *B. alpinus/polaris* | 0.667 |
| *B. pascuorum* | *B. balteatus* | 0.617 |
| *B. pascuorum* | *B. bohemicus* | 0.732 |
| *B. pascuorum* | *B. hortorum* | 0.719 |
| *B. pascuorum* | *B. jonellus* | 0.509 |
| *B. pascuorum* | *B. lapponicus* | 0.828 |
| *B. pascuorum* | *B. lucorum* | 0.568 |
| *B. pascuorum* | *B. monticola* | 0.771 |
| *B. pratorum* | *B. alpinus/polaris* | 0.442 |
| *B. pratorum* | *B. balteatus* | 0.450 |
| *B. pratorum* | *B. bohemicus* | 0.350 |
| *B. pratorum* | *B. hortorum* | 0.635 |
| *B. pratorum* | *B. jonellus* | 0.744 |
| *B. pratorum* | *B. lapponicus* | 0.697 |
| *B. pratorum* | *B. lucorum* | 0.374 |
| *B. pratorum* | *B. monticola* | 0.638 |
| *B. pratorum* | *B. pascuorum* | 0.599 |

| Table 10: **Summary of an ordinary least squares model investigating the change in bumblebee (*Bombus* spp.) intertegular distance over the course of the sampling season, when considering all individuals.** Values in bold were considered statistically significant (*p* ≤0.05). Multiple R^2^ = 0.351. *N* = 709. | | | | |
| --- | --- | --- | --- | --- |
| Coefficient | Estimate | Standard error | *t* value | *p* value |
| Intercept | 875 | 44.5 | 19.7 | **<0.001** |
| Date | -0.0491 | 0.00251 | -19.6 | **<0.001** |

| Table 11: **Summary of a linear mixed-effects model investigating the change in bumblebee (*Bombus* spp.) intertegular distance over the course of the sampling season, when considering all individuals.** Bumblebee species were included as random intercepts. Values in bold were considered statistically significant (*p* ≤0.05). Marginal R^2^ = 0.314, conditional R^2^ = 0.541. *N* = 709. | | | | |
| --- | --- | --- | --- | --- |
| Coefficient | Estimate | Standard error | *t* value | *p* value |
| Intercept | 838 | 39.6 | 21.1 | **<0.001** |
| Date | -0.0470 | 0.00225 | -20.9 | **<0.001** |
| Random intercepts | Variance | Standard deviation |  |  |
| *Bombus* species | 0.355 | 0.596 |  |  |
| Residual | 0.0718 | 0.847 |  |  |

| Table 12: **Summary of a linear mixed-effects model investigating the change in bumblebee (*Bombus* spp.) intertegular distance over the course of the sampling season, when considering all individuals.** Bumblebee species had random intercepts and gradients (date). Values in bold were considered statistically significant (*p* ≤0.05). Marginal R^2^ = 0.274, conditional R^2^ = 0.602. *N* = 709. | | | | |
| --- | --- | --- | --- | --- |
| Coefficient | Estimate | Standard error | *t* value | *p* value |
| Intercept | 839 | 39.7 | 21.1 | **<0.001** |
| Date | -0.0471 | 0.00224 | -21.0 | **<0.001** |
| Random effects | Variance | Standard deviation |  |  |
| *Bombus* species (Intercepts) | 0.713 | 0.845 |  |  |
| Date (Gradients) | <0.001 | <0.001 |  |  |
| Residual | 0.713 | 0.845 |  |  |

| Table 13: **Summary of an ordinary least squares model investigating the change in bumblebee (*Bombus* spp.) intertegular distance with increasing elevation, when considering all individuals.** Values in bold were considered statistically significant (*p* ≤0.05). Multiple R^2^ = 0.0258. *N* = 709. | | | | |
| --- | --- | --- | --- | --- |
| Coefficient | Estimate | Standard error | *t* value | *p* value |
| Intercept | 4.26 | 0.189 | 22.5 | **<0.001** |
| Altitude | 0.00121 | <0.001 | 4.33 | **<0.001** |

| Table 14: **Summary of a linear mixed-effects model investigating the change in bumblebee (*Bombus* spp.) intertegular distance with increasing elevation, when considering all individuals.** Bumblebee species were included as random intercepts. Values in bold were considered statistically significant (*p* ≤0.05). Marginal R^2^ <0.001, conditional R^2^ = 0.241. *N* = 709. | | | | |
| --- | --- | --- | --- | --- |
| Coefficient | Estimate | Standard error | *t* value | *p* value |
| Intercept | 5.34 | 0.262 | 20.4 | **<0.001** |
| Altitude | <0.001 | 0.282 | 0.563 | **0.574** |
| Random intercepts | Variance | Standard deviation |  |  |
| *Bombus* species | 0.370 | 0.608 |  |  |
| Residual | 1.17 | 1.08 |  |  |
| Table 15: **Summary of a linear mixed-effects model investigating the change in bumblebee (*Bombus* spp.) intertegular distance with increasing elevation, when considering all individuals.** Bumblebee species had random intercepts and gradients (elevation). Values in bold were considered statistically significant (*p* ≤0.05). Marginal R^2^ = 0.00260, conditional R^2^ = 0.256. *N* = 709. | | | | |
| Coefficient | Estimate | Standard error | *t* value | *p* value |
| Intercept | 5.17 | 0.216 | 23.9 | **<0.001** |
| Altitude | <0.001 | <0.001 | 1.13 | 0.278 |
| Random effects | Variance | Standard deviation |  |  |
| *Bombus* species (Intercepts) | 0.0955 | 0.309 |  |  |
| Altitude (Gradients) | <0.001 | <0.001 |  |  |
| Residual | 1.17 | 1.08 |  |  |

| Supplementary Table 16: **Species turnover over space and time using Bray-Curtis distances.** Bray-Curtis distance is broken down into its two additive components (sensu Balsega, 2013): balanced variation in abundance (where bees of one species at one site are replace by the same number of bees of another species at another site; D_BC-BAL_) and abundance gradients (where sites differ in the number of bees; D_BC-GRA_). We only considered bees observed foraging and with a measurement of intertegular distance. | | | |
| --- | --- | --- | --- |
| Bray Curtis | | | |
|  | Early season, low elevation | Early season, high elevation | late season, low elevation |
| Early season, high elevation | 0.533 |  |  |
| late season, low elevation | 0.474 | 0.781 |  |
| Late season, high elevation | 0.322 | 0.672 | 0.440 |
| D_BC-BAL_ | | | |
|  | Early season, low elevation | Early season, high elevation | late season, low elevation |
| Early season, high elevation | 0.125 |  |  |
| late season, low elevation | 0.0682 | 0.125 |  |
| Late season, high elevation | 0.0795 | 0.0625 | 0.305 |
| D_BC-GRA_ | | | |
|  | Early season, low elevation | Early season, high elevation | late season, low elevation |
| Early season, high elevation | 0.408 |  |  |
| late season, low elevation | 0.406 | 0.656 |  |
| Late season, high elevation | 0.243 | 0.610 | 0.135 |

| Supplementary Table 17: **Constrained-functional-group turnover over space and time using Bray-Curtis distances.** Bray-Curtis distance is broken down into its two additive components (sensu Balsega, 2013): balanced variation in abundance (where bees of one functional group at one site are replace by the same number of bees of another functional group at another site; D_BC-BAL_) and abundance gradients (where sites differ in the number of bees; D_BC-GRA_). We only considered bees observed foraging and with a measurement of intertegular distance. | | | |
| --- | --- | --- | --- |
| Bray Curtis | | | |
|  | Early season, low elevation | Early season, high elevation | late season, low elevation |
| Early season, high elevation | 0.500 |  |  |
| late season, low elevation | 0.776 | 0.945 |  |
| Late season, high elevation | 0.623 | 0.792 | 0.269 |
| D_BC-BAL_ | | | |
|  | Early season, low elevation | Early season, high elevation | late season, low elevation |
| Early season, high elevation | 0.0625 |  |  |
| late season, low elevation | 0.602 | 0.781 |  |
| Late season, high elevation | 0.489 | 0.406 | 0.0927 |
| D_BC-GRA_ | | | |
|  | Early season, low elevation | Early season, high elevation | late season, low elevation |
| Early season, high elevation | 0.438 |  |  |
| late season, low elevation | 0.173 | 0.164 |  |
| Late season, high elevation | 0.135 | 0.386 | 0.177 |

| Table 18: **Turnover over space and time using Jaccard dissimilarity.** We only considered bees observed foraging and with a measurement of intertegular distance. | | | |
| --- | --- | --- | --- |
| Species turnover | | | |
|  | Early season, low elevation | Early season, high elevation | late season, low elevation |
| Early season, high elevation | 0.333 |  |  |
| late season, low elevation | 0.154 | 0.333 |  |
| Late season, high elevation | 0.231 | 0.417 | 0.0833 |
| Functional turnover (constrained clusters) | | | |
|  | Early season, low elevation | Early season, high elevation | late season, low elevation |
| Early season, high elevation | 0.455 |  |  |
| late season, low elevation | 0.417 | 0.615 |  |
| Late season, high elevation | 0.250 | 0.462 | 0.167 |

| Supplementary Table 19: **Summary of network-level indices.** *Z*- and *p*-values (in parentheses, respectively) were calculated by comparing the indices for each network with the distributions of the indices generated using null network (using “vaznull” null networks from “bipartite” package; Dormann et al., 2009). Values in bold were considered statistically significant (*p* <0.05). | | | | | | |
| --- | --- | --- | --- | --- | --- | --- |
|  | Network nodes | Bee species or functional group richness | Plant species richness | Number of interactions | Weighted connectance | Modularity |
| Early season, low elevation | Species | 12 | 6 | 88 | 0.245 (-1.25, 0.211) | 0.154 (0.644, 0.519) |
|  | Functional groups | 9 | 6 | 88 | 0.285 (0.961, 0.337) | 0.116 (-0.432, 0.666) |
| Late season, low elevation | Species | 12 | 16 | 224 | **0.147 (-3.82, <0.001)** | **0.241 (5.17, <0.001)** |
|  | Functional groups | 10 | 16 | 224 | 0.170 (-1.92, 0.0551) | 0.136 (0.228, 0.820) |
| Early season, high elevation | Species | 8 | 6 | 32 | 0.189 (-1.62, 0.106) | 0.303 (-0.342, 0.973) |
|  | Functional groups | 8 | 6 | 32 | 0.203 (-0.998, 0.319) | 0.346 (0.685, 0.493) |
| Late season, high elevation | Species | 11 | 21 | 151 | **0.168 (-2.02, 0.0434)** | **0.334 (3.98, <0.001)** |
|  | Functional group | 12 | 21 | 151 | **0.147 (-3.04, 0.00238)** | 0.245 (1.06, 0.295) |

| Supplementary Table 20: **Generalised least squares models summarising the change in weighted connectance, weighted quantitative modularity and number of remaining flower species as constrained functional groups are randomly and sequentially removed from species nodes in networks across different points in space and time.** Relationships are based on 1,000 iterations of the random removal of functional groups. An exponent of the variance covariate was included in each model to account for the heterogeneity of variance in the residuals; the exception to this was weighted quantitative modularity, due to problems with model convergence. S.E. = standard error of model estimate; AUC = area under the curve. AUC was standardised by dividing the raw value by the product of the number of functional groups removed from species and the maximum value of the response variable. Values in bold are statistically significant (*p* <0.05). | | | | | | | | |
| --- | --- | --- | --- | --- | --- | --- | --- | --- |
| Weighted connectance | | | | | | | | |
|  | Intercept  (±S.E.) | Intercept  *t-* & *p*-value | β  (±S.E.) | β  *t-* & *p*-value | β^2^  (±S.E.) | β^2^  *t-* & *p*-value | Raw AUC | Standardised AUC |
| Early season, low elevation | 0.248  (±<0.001) | **991,**  **<0.001** | 0.00105  (±0.001) | **25.6, <0.001** | <0.001  (±<0.001) | **11.3, <0.001** | 11.1 | 0.885 |
| Late season, low elevation | 0.150  (±<0.001) | **1,320,**  **<0.001** | <-0.001  (±<0.001) | **-6.65,**  **<0.001** | <0.001  (±<0.001) | **81.1,**  **<0.001** | 6.38 | 0.732 |
| Early season, high elevation | 0.191  (±<0.001) | **230,**  **<0.001** | 0.00646  (±<0.001) | **15.1,**  **<0.001** | <0.001  (±<0.001) | **13.0,**  **<0.001** | 3.08 | 0.738 |
| Late season, high elevation | 0.173  (±<0.001) | **1,080,**  **<0.001** | <-0.001  (±<0.001) | **-3.64,**  **<0.001** | <0.001  (±<0.001) | **-106,**  **<0.001** | 8.72 | 0.764 |
| Remaining flower species | | | | | | | | |
|  | Intercept  (±S.E.) | Intercept  *t-* & *p*-value | β  (±S.E.) | β  *t-* & *p*-value | β^2^  (±S.E.) | β^2^  *t-* & *p*-value | Raw AUC | Standardised AUC |
| Early season, low elevation | 6.01  (±0.00908) | **662,**  **<0.001** | -0.0507  (±0.00125) | **-40.6,**  **<0.001** | <-0.001  (±<0.001) | **-25.6,**  **<0.001** | 182 | 0.755 |
| Late season, low elevation | 15.8  (±0.0102) | **1,540,**  **<0.001** | 0.00304  (±0.00193) | 1.58,  0.114 | 0.00680  (±<0.001) | **-101,**  **<0.001** | 458 | 0.828 |
| Early season, high elevation | 5.97  (±0.0227) | **263,**  **<0.001** | -0.219  (±0.00911) | **-24.1,**  **<0.001** | -0.00514  (±<0.001) | **-6.84,**  **<0.001** | 52.9 | 0.738 |
| Late season, high elevation | 20.7  (±0.0171) | **1,210,**  **<0.001** | -0.0516  (±0.00234) | **-22.1,**  **<0.001** | -0.00652  (±<0.001) | **-106,**  **<0.001** | 671 | 0.753 |
| Weighted quantitative modularity | | | | | | | | |
|  | Intercept  (±S.E.) | Intercept  *t-* & *p*-value | β  (±S.E.) | β  *t-* & *p*-value | β^2^  (±S.E.) | β^2^  *t-* & *p*-value | Raw AUC | Standardised AUC |
| Early season, low elevation | 0.164  (±<0.001) | **194,**  **<0.001** | -0.00204  (±<0.001) | **-20.4,**  **<0.001** | <0.001  (±<0.001) | **29.1, <0.001** | 6.49 | 0.816 |
| Late season, low elevation | 0.238  (±<0.001) | **255,**  **<0.001** | <0.001  (±0.001) | **6.73,**  **<0.001** | <-0.001  (±<0.001) | **-21.6,**  **<0.001** | 7.80 | 0.926 |
| Early season, high elevation | 0.314  (±0.00326) | **96.2,**  **<0.001** | --0.00746  (±0.00119) | **-6.26,**  **<0.001** | <0.001  (±<0.001) | **5.39,**  **<0.001** | 3.52 | 0.934 |
| Late season, high elevation | 0.323  (±<0.001) | **357,**  **<0.001** | 0.00279  (±<0.001) | **28.8,**  **<0.001** | <-0.001  (±0.001) | **-23.5,**  **<0.001** | 15.1 | 0.973 |

| Supplementary Table 21: **Generalised least squares models summarising the change in weighted connectance, number of remaining flower species and weighted quantitative as species are randomly and sequentially removed from functional-group nodes in networks across different points in space and time.** Relationships are based on 1,000 iterations of the random species removal. An exponent of the variance covariate was included in each model to account for the heterogeneity of variance in the residuals; the exception to this was weighted quantitative modularity, due to problems with model convergence. S.E. = standard error of model estimate; AUC = area under the curve. AUC was standardised by dividing the raw value by the product of the number of species removed from functional groups and the maximum values of the response variable. Values in bold are statistically significant (*p* <0.05). | | | | | | | | |
| --- | --- | --- | --- | --- | --- | --- | --- | --- |
| Weighted connectance | | | | | | | | |
|  | Intercept  (±S.E.) | Intercept  *t-* & *p*-value | β  (±S.E.) | β  *t-* & *p*-value | β^2^  (±S.E.) | β^2^  *t-* & *p*-value | Raw AUC | Standardised AUC |
| Early season, low elevation | 0.287  (±<0.001) | **1,230,**  **<0.001** | -0.00165  (±<0.001) | **-42.8,**  **<0.001** | <0.001  (±<0.001) | **54.6,**  **<0.001** | 11.6 | 0.887 |
| Late season, low elevation | 0.172  (±<0.001) | **1,170,**  **<0.001** | 0.00192  (±<0.001) | **58.1,**  **<0.001** | <0.001  (±<0.001) | **21.7,**  **<0.001** | 7.61 | 0.791 |
| Early season, high elevation | 0.196  (±<0.001) | **275,**  **<0.001** | 0.00144  (±<0.001) | **4.02,**  **<0.001** | <0.001  (±<0.001) | **20.8,**  **<0.001** | 2.87 | 0.756 |
| Late season, high elevation | 0.151  (±<0.001) | **1,130,**  **<0.001** | <0.001  (±<0.001) | **14.9,**  **<0.001** | <0.001  (±<0.001) | **92.5,**  **<0.001** | 8.55 | 0.692 |
| Remaining flower species | | | | | | | | |
|  | Intercept  (±S.E.) | Intercept  *t-* & *p*-value | β  (±S.E.) | β  *t-* & *p*-value | β^2^  (±S.E.) | β^2^  *t-* & *p*-value | Raw AUC | Standardised AUC |
| Early season, low elevation | 5.92  (±0.00871) | **680,**  **<0.001** | -0.0234  (±0.00120) | -19.4,  <0.001 | -0.00145  (±<0.001) | **-43.8,**  **<0.001** | 187 | 0.791 |
| Late season, low elevation | 15.8  (±0.00914) | **1,730,**  **<0.001** | 0.0350  (±0.00185) | **18.9,**  **<0.001** | -0.00771  (±<0.001) | **-112,**  **<0.001** | 465 | 0.838 |
| Early season, high elevation | 5.88  (±0.0199) | **295,**  **<0.001** | -0.0623  (±0.00829) | -7.52,  <0.001 | -0.0161  (±<0.001) | **-22.9,**  **<0.001** | 56.7 | 0.804 |
| Late season, high elevation | 20.5  (±0.0164) | **1,250,**  **<0.001** | -0.0773  (±0.00225) | -34.3,  <0.001 | -0.00511  (±<0.001) | **-85.7,**  **<0.001** | 675 | 0.765 |
| Weighted quantitative modularity | | | | | | | | |
|  | Intercept  (±S.E.) | Intercept  *t-* & *p*-value | β  (±S.E.) | β  *t-* & *p*-value | β^2^  (±S.E.) | β^2^  *t-* & *p*-value | Raw AUC | Standardised AUC |
| Early season, low elevation | 0.124  (±<0.001) | **166,**  **<0.001** | <-0.001  (±0.001) | -11.2, <0.001 | <0.001  (±<0.001) | **27.7,**  **<0.001** | 5.47 | 0.752 |
| Late season, low elevation | 0.134  (±<0.001) | **158,**  **<0.001** | <0.001  (±<0.001) | 3.78,  <0.001 | <0.001  (±<0.001) | **10,**  **<0.001** | 5.39 | 0.823 |
| Early season, high elevation | 0.358  (±0.00318) | **113,**  **<0.001** | -0.00249  (±0.00116) | -2.14,  0.0327 | <-0.001  (±<0.001) | **-1.97,**  **0.0484** | 4.01 | 0.934 |
| Late season, high elevation | 0.256  (±<0.001) | **331,**  **<0.001** | -0.00123  (±<0.001) | -14.6,  <0.001 | <0.001  (±<0.001) | **9.32,**  **<0.001** | 10.3 | 0.940 |

**Bibliography:**

Bates, D., Maechler, M., Bolker, B., Walker, S., Christensen, R.H.B., Singmann, H., *et al.* (2018). Package ‘lme4.’ *Version*, 1, 437.

Cox, T. (2018). Topology of plant-bumblebee interaction networks along an elevational gradient in the Arctic. Imperial College London, UK.

Dormann, C.F., Frund, J., Bluthgen, N. & Gruber, B. (2009). Indices, Graphs and Null Models: Analyzing Bipartite Ecological Networks. *Open Ecol. J.*, 2, 7–24.

Martin, S.J., Carruthers, J.M., Williams, P.H. & Drijfhout, F.P. (2010). Host specific social parasites (Psithyrus) indicate chemical recognition system in bumblebees. *J. Chem. Ecol.*, 36, 855–863.
